# Mechanochemical feedback between confinement and actin crosslinking drives the shape dynamics of liquid-like droplets

**DOI:** 10.1101/2025.06.13.659650

**Authors:** Daniel Mansour, Dominique Jordan, Caleb Walker, Aravind Chandrasekaran, Christopher T. Lee, Kristin Graham, Jeanne Stachowiak, Padmini Rangamani

## Abstract

Several actin-binding proteins can form liquid-liquid phase-separated condensates that promote actin filament assembly and bundling, which is crucial for local actin network organization. Previous studies have established that phase-separated condensates composed of actin-binding proteins, such as vasodilator-stimulated phosphoprotein (VASP) and Lamellipodin (Lpd), restrict the organization of actin filaments to structures such as rings, shells, discs, and rods through kinetic trapping. However, the mechanism by which crosslinker multivalency, actin growth, and condensate properties tune actin organization and droplet shape is not well understood. Using a combination of agent-based simulations and experiments, we find that the deformability of the droplet interface allows for the emergence of not just tightly-bundled actin rings but also weakly-bundled actin discs. We find two major quantitative relationships between actin bundling and droplet deformation. The first relationship shows that the crosslinked bundle thickness and droplet diameter followed a power law, consistent with experiments. The second one is that the kinetics of droplet deformation follows a dynamic snapping behavior that depends on the droplet surface tension and the multivalent VASP-actin binding kinetics. We predicted that these two relationships were generalizable to dynamic multimers and to weak actin crosslinkers. Our predictions were experimentally tested using two additional condensate-forming proteins, lamellipodin and RGG. Taken together, we show that mechanochemical feedback between the droplet interface properties and crosslinker multivalency tune actin organization and control the dynamics of droplet deformation by actin networks.

## Introduction

The interaction between actin filaments and actin-binding proteins (ABPs) leads to the formation of a variety of actin network architectures^1^. There are over 160 ABPs, and each of them has specific actin-binding roles that transform unorganized actin monomers and filaments into specialized structures that drive complex biological processes^2,3^. Despite the diversity in ABPs, their effect on actin filaments can be broadly classified as tuning of filament assembly and disassembly rates, branching, severing and annealing, and bundling. Reconstituted actin networks under confinement have been pivotal in highlighting the principles of actin organization^4^. Similar observations of actin ring formation have been observed in both experimental studies with GUVs^5^ and vesicles^6^.

Recent discoveries have shown that several ABPs can form condensates through the process of liquid-liquid phase-separation, both in vitro and in vivo^7–16^. We have established how the spatial confinement of ABPs through phase separation affects actin filament assembly and bundling^8,9,17,18^. For example, we recently showed that VASP forms phase-separated droplets, and in such droplets, monomeric actin can assemble into filamentous actin networks of different configurations^9^. Depending on the ratio of actin to VASP, the resulting actin networks can form shells, rings, or ellipsoidal discs^9,18^. Furthermore, the formation of bundled actin networks (rings) was a precursor to droplet deformation, where spherical VASP droplets transitioned to ellipsoidal droplets and finally rods^9,18^. More recently, we also showed experimentally that such actin bundling did not require native tetrameric strong bundling proteins such as VASP but was also possible due to proteins such as lamellipodin (Lpd), which bind actin but lack inherent polymerase activity^19–22^. Additionally, we showed that when Eps15, a condensate-forming protein without any known interaction with actin, is fused with Lifeact, a general F-actin binding motif, the resulting Eps15-Lifeact composite promotes actin filament assembly and bundling^8^. Using an agent-based modeling approach, we previously investigated how interactions between VASP and actin in rigid, non-deformable droplets can lead to the formation of rings versus shells^18^. We showed that rings of actin resulted from kinetic trapping, where for a fixed filament growth rate, the binding and unbinding rate of VASP to actin allowed for a sufficient residence time of VASP on one actin filament such that another filament could bind and zipper^9^. Subsequently, we also showed that dynamic dimers and multimers, representing Lpd and Eps15-Lifeact composites, also demonstrated rings versus shells resulting from kinetic trapping^8^. In the case of GUV-encapsulated actin, membrane-actin tethering has been shown to be critical for ring formation^6^. These experimental observations suggest that there are general principles underlying the assembly of actin shells, rings, and discs in protein droplets.

However two questions remain: What is the role of mechanical feedback from deformable boundary confinement in tuning actin organization within the droplet? How do the specific crosslinker biochemical properties coordinate droplet deformation? To address these, we used a combination of computational and experimental approaches. We modeled the ABP droplet using dynamically deformable ellipsoidal boundaries and represented both the ABP-ABP and ABP-actin kinetic interactions explicitly (**Fig. 1**). Our simulations revealed that the condensate environment tunes the nature of actin organization through kinetic trapping, thereby controlling droplet shape. We predicted that the condensate diameter has a power law scaling relationship with the actin bundle thickness required to deform the spherical droplet. These predictions were tested using experiments. We also showed in simulations and experiments that capping protein tunes the filament length distribution and can alter the onset and extent of VASP droplet deformation. Finally, we showed that the temporal behavior of droplet deformation has a dynamic snapping behavior depending on the droplet mechanical properties and the crosslinker kinetic properties. Thus, our study produces a comprehensive, general framework to understand the relationship between droplet deformation and the mechanochemical environment of the condensate.

**Figure 1:**
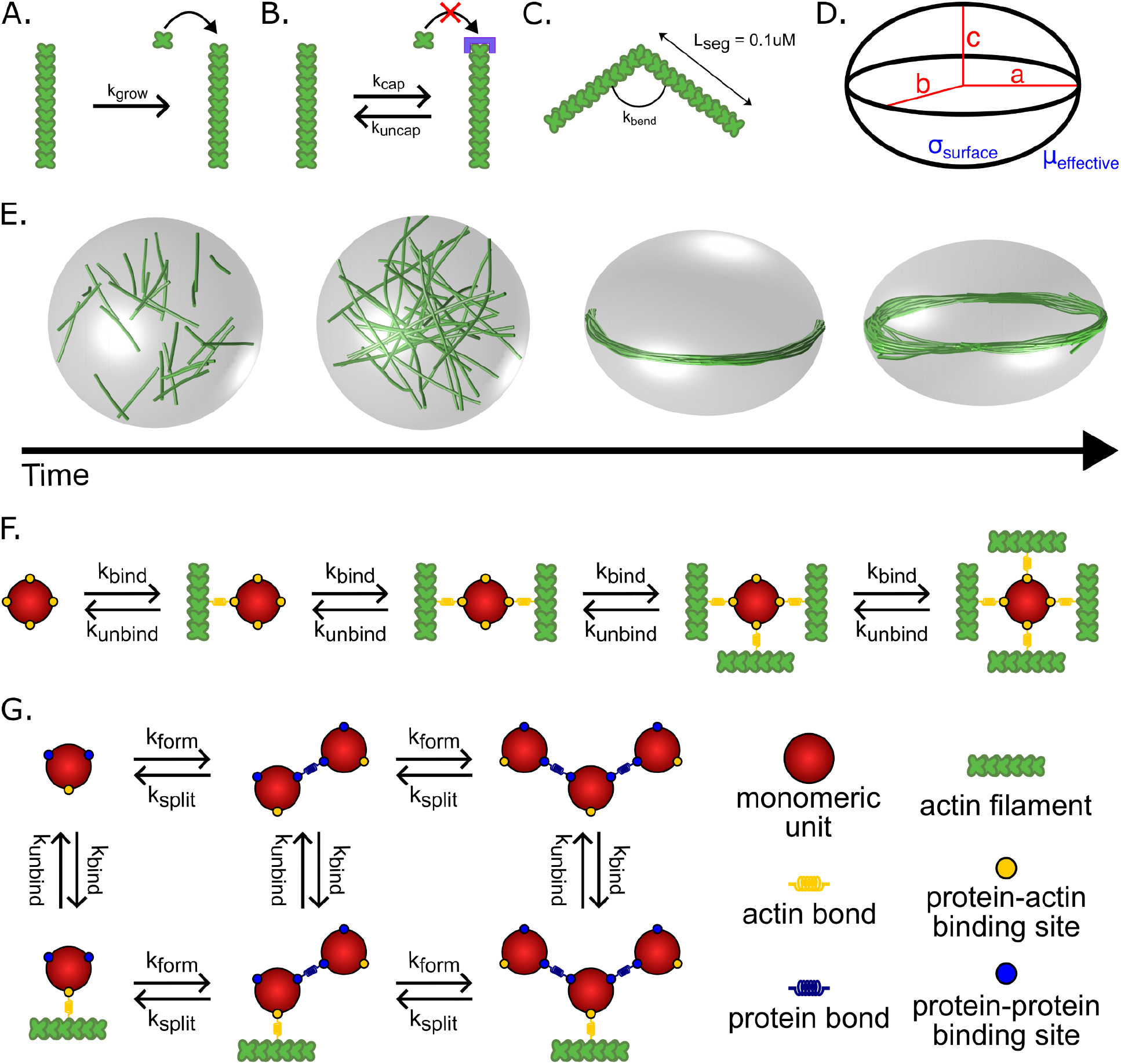
Simulation framework for capturing actin network properties in droplets of actin-binding proteins (ABPs). **A)** Actin filaments are modeled with a deterministic growth rate k_grow_ that was chosen to allow the filaments to grow to the length of the circumference of the starting spherical droplet space by the end of the simulation (600 s). The growth rate is set at k_grow_ = 0.0103 μm/s unless otherwise stated. **B)** Capping protein is implicitly modeled using rates of capping and uncapping that are specified at the plus (+) ends of the actin filaments. Capped filaments do not participate in filament growth. **C)** Actin filaments are modeled as inextensible segments of length L_seg_ that can bend along hinge points based on flexural rigidity k_bend_. **D)** In the dynamic ellipsoid model, the geometry of the condensate is constrained to an ellipsoid with principal axes a, b, and c. **E)** Condensate boundary deformation is regulated by the condensate surface tension σ_surface_ and the effective viscosity µ_effective_. The deformation of the condensate is determined by the balance between the condensate surface tension σ_surface_ and the point forces exerted by the filaments on the boundary. The time evolution of deformation is further modulated by the effective viscosity of the droplet, µ_effective_, essentially a damping parameter, which slows the deformation of the axes over time. **F)** Schematic of the different combinations of possible binding and unbinding reactions that tetrameric crosslinkers can undergo. Each of the four actin-binding domains in the tetrameric crosslinker binds actin with rate k_bind_ and unbinds in a force-sensitive manner with unbinding rate k_unbind_. **G)** Schematic depicting the various actin-binding and protein-protein binding reactions that monomeric units in the dynamic multimerization model can undergo. Each monomeric unit has a single actin-binding site and two protein-protein binding sites to facilitate the formation of large multimers. For each protein-protein binding domain, monomers bind to each other with a multimer formation rate k_form_ and split in a force-sensitive manner with a multimer splitting rate k_split_.

## Methods

### Model development

We constructed an agent-based model in Cytosim to investigate the interplay between the physical and chemical properties of the ABP droplet and the actin network. Cytosim simulates the chemical dynamics and mechanical properties of cytoskeletal filament networks^23^. This modeling framework allows us to operate explicitly with three distinct components: filaments, the condensate space, and crosslinkers.

Actin filaments are represented as inextensible fibers that elongate at a constant rate to reflect polymerization (**Fig. 1A**). We assume that only the plus end elongates while the minus end is stable. These assumptions are consistent with the role of VASP as a processive polymerase that mediates up to a three-fold increase in elongation rate at bound barbed ends compared to free barbed ends^24^, and based on previous models of actin filament elongation with a net elongation at the plus ends^18^. Additionally, in this study, we introduce a method for implicitly modeling the effect of capping protein by stochastically stopping (capped state) and restarting (uncapped state) elongation via a capping “reaction” governed by the rate parameters k_cap_ and k_uncap_ (**Fig. 1B**). Finally, actin filaments are composed of a series of linear segments of length 0.1 µm that are connected by hinge points to allow for bending (**Fig. 1C**). Cytosim computes the bending energy of the fiber using the specified flexural rigidity k_bend_ in the input parameters^25^.

The condensate space is the defined 3D geometry where reactions, filament dynamics, and diffusion occur as before^8,17,18^. Whereas our previous models used a spherical geometry that was rigidly enforced or deformed in a quasi-static manner^8,17,18^, we now incorporate a deformable ellipsoid geometry that dynamically adjusts the space and principal axes with each timestep in accordance with the forces acting on the boundary against the surface tension of the droplet (**Fig. 1D-E**). This framework for Cytosim was developed in Dmitrieff et al. 2017^26^ to understand the role of cortical tension in microtubule assembly. Here, we use the framework to represent the deformable boundary of condensates. By modeling the condensate geometry as a dynamic ellipsoid, we investigated the feedback between the condensate properties and the actin network. Deformation is calculated by the balance between the interfacial surface tension and the point forces exerted by the filaments on the boundary and is subject to volume incompressibility constraints^26^. The speed at which deformation occurs, but not the final shape, is further modulated by an effective viscosity, µ_effective_, which acts as a damping parameter to slow the deformation of the ellipse axes over time^26^.

The ABPs are represented as spherical solids with a radius of 30 nm and can be modeled with two different kinds of binding sites: sites that bind actin to promote bundling and sites that bind to other ABP molecules to promote multimerization. Each binding site has a specified binding rate constant k_bind_ and binding distance [30 nm] that specifies the radius within which targets are considered in the binding reaction^18^. Additionally, the unbinding rate k_unbind_ specifies the rate constant used in a force-dependent kinetics (Bell’s law model) representation of slip bond unbinding kinetics^27,28^. We investigated the role of two distinct types of actin crosslinkers: static tetrameric actin crosslinkers representing VASP^17,18^ and dynamically multimerizing actin crosslinkers representing the transient interactions that may occur between monomeric ABPs in an inherently multivalent condensate environment^8^. As a tetrameric actin crosslinker, VASP is modeled with four actin-binding sites that enable it to undergo a series of reversible reactions that allow each VASP molecule to exist in five different states depending on the number of bound actin filaments (**Fig. 1F**)^18^. Additionally, we introduced modifications to the Cytosim codebase to model dynamically multimerizing monomers based on a specified multimer formation rate k_form_ and splitting rate k_split_, as first introduced in Walker et al. 2025^8^. These dynamic crosslinkers differ from static crosslinkers in that each crosslinker molecule is a monomer modeled with a single actin-binding site and two monomer-monomer binding sites to allow each monomer to bind with up to two other monomers to dynamically form multimers such as dimers, trimers, tetramers, etc. The inclusion of the monomer-monomer binding sites extends the framework to allow for multimeric crosslinker formation. Due to the large number of possible states, we categorize each monomer unit by the number of other monomers it is bound to [0, 1, 2] and by whether or not it is bound to an actin filament [B] or free [F], thus resulting in six different states that each monomeric unit can cycle between through a series of actin or monomer binding and unbinding reactions (**Fig. 1G**). Specific parameters used to set up the Cytosim model can be found in **Supplementary Table S1** while the list of parameters varied in each set of simulations can be found in **Supplementary Table S2**.

The Cytosim simulations were performed on the Triton Shared Computing Cluster (TSCC) at the San Diego Supercomputer Center (SDSC)^29^. Due to the stochastic nature of the interactions in our simulations, 10 replicate simulations were performed per condition to yield a sufficiently large sample size for statistical analyses of the results. All Cytosim trajectories were initialized with a random distribution of filaments and crosslinker species. All data analysis for simulations were performed on Python 3.11.5. The custom Cytosim source code used to generate trajectories and the Python scripts used to analyze trajectories will be made available on publication.

### Experimental methods

Condensates composed of VASP, monomer mini-Lpd, or RGG were formed as described previously with minor alterations^8,9^. Briefly, the given concentrations of protein (see text) were mixed with 3% (w/v) PEG-8000 in 20 mM Tris pH 7.4, 5 mM TCEP, and 150 mM NaCl for VASP condensates or 50 mM NaCl for RGG and monomer mini-Lpd condensates. PEG was added last to induce condensate formation after the protein was well-mixed in the solution. For the actin assembly assay, condensates were formed for ten minutes (with time starting after PEG addition) prior to addition of G-actin. Following actin addition the sample was incubated at room temperature for 15 minutes prior to imaging to allow for actin polymerization. Samples were prepared for microscopy in 3.5 mm diameter wells formed using biopsy punches to create holes in 1.6 mm thick silicone gaskets (Grace Biolabs) on Hellmanex III cleaned, no. 1.5 glass coverslips (VWR). Coverslips were passivated using poly-L-lysine conjugated PEG chains (PLL-PEG). After addition of samples to these wells, a second coverslip was placed on top to seal the imaging chamber against evaporation during imaging. Fluorescence microscopy was performed using an Olympus SpinSR10-Yokogawa SoRa spinning disk confocal microscope with a Hamamatsu Orca Flash 4.0V3 Scientific CMOS camera.

ImageJ was used to quantify the distribution of condensate characteristics. Specifically, condensates were selected using thresholding in the brightest channel and shape descriptors (i.e., diameter, aspect ratio, etc.), and protein fluorescent intensities were measured using the built-in analyze particles function. For aspect ratio analysis condensates that had come into contact with other condensates were removed from the analysis to avoid any skewing of data from misrepresentation of single condensate deformation.

Detailed methods for the proteins and experimental analysis can be found in the **Supplemental Methods**.

## Results

Our investigations are organized as follows: we first show that in droplets containing VASP, allowing for droplet deformation leads to the emergence of discs (weakly-bundled actin structures) that are oriented along the axis of deformation. We found that a power law relationship exists between the initial radius of the droplet and the number of filaments for a particular aspect ratio. Our simulations reveal that the kinetics of droplet deformation follows a dynamic snapping behavior, a snap-through of the shape of the droplet that occurs over time, depending on the organization of the actin network in the VASP droplet. We then predicted that actin filament length plays an important role in determining the onset and extent of droplet deformation; this prediction is verified by experimentally changing the capping protein concentration. We also show that all our observations are generalizable to ABPs that form dynamic multimers, establishing that actin bundling and droplet deformation do not require strong bundling proteins. Even in the absence of a specific ABP in the droplet, we show that actin filaments will form discs that align along the axis of deformation, and therefore, concentrating actin in phase-separated containers is a sufficient condition for bundling and force generation. Finally, we show that the biochemical properties of actin crosslinkers drive actin structure formation by influencing the interplay between droplet shape and bundle alignment.

### Deformable VASP condensates produce tightly bundled actin rings and weakly bundled discs

In our previous work, we simulated actin bundling in rigid VASP droplets^8,17,18^. However, in the experiments, we observe droplet deformation^8,9,17,18^. Here, we first relaxed the assumption of the rigid droplet and asked whether the deformability of the droplet influences actin assembly. In our simulations, the droplet is allowed to deform with a given surface tension and a viscous dissipation parameter along with a constant volume constraint^26^. To fully map the interaction between the droplet interface and the actin bundling, we simulated a range of binding and unbinding kinetics of VASP with actin filaments in deformable droplets (**Fig. 2**). In these deformable droplets, we observed the formation of weakly-bundled discs of actin filaments and rings; the weakly-bundled discs were seen in the regime of weak VASP binding kinetics (*k* _*bind*_ ≥ 0. 1 and *k*_*bind*_ < *k*_*unbind*_ or *k*_*bind*_ < 0. 1) (**Fig. 2A**). This is different from the case of rigid droplets, where we observe shells and rings (**Fig. S1** and Chandrasekaran et al.^18^). The disc-shaped bundles that emerge in deformable droplets appeared to align themselves along the axes of droplet deformation to minimize the energetic penalty to filament bending (**Fig. 2A**). We quantified the thickness of the discs by analyzing the surface area fraction of the droplet covered by actin^18^. The differences in surface area fraction for rings and discs are shown in **Fig. 2A**, where rings form with a low surface area fraction, while discs have a much larger surface area fraction similar to shells seen in rigid droplets (**Fig. 2B, Fig. S1**). Additionally, when comparing the fraction of VASP tetramers bound to actin filaments, almost no VASP tetramers were bound to any actin filaments in simulations with kinetic conditions that favored disc formation (**Fig. 2C**). This is in contrast to rings and shells, where kinetic trapping by VASP tetramers was required to tightly bundle actin filaments into rings or tightly confine the actin filaments within the original spherical geometry of the droplet so that the filament network develops into shell-like shapes^18^.

**Figure 2:**
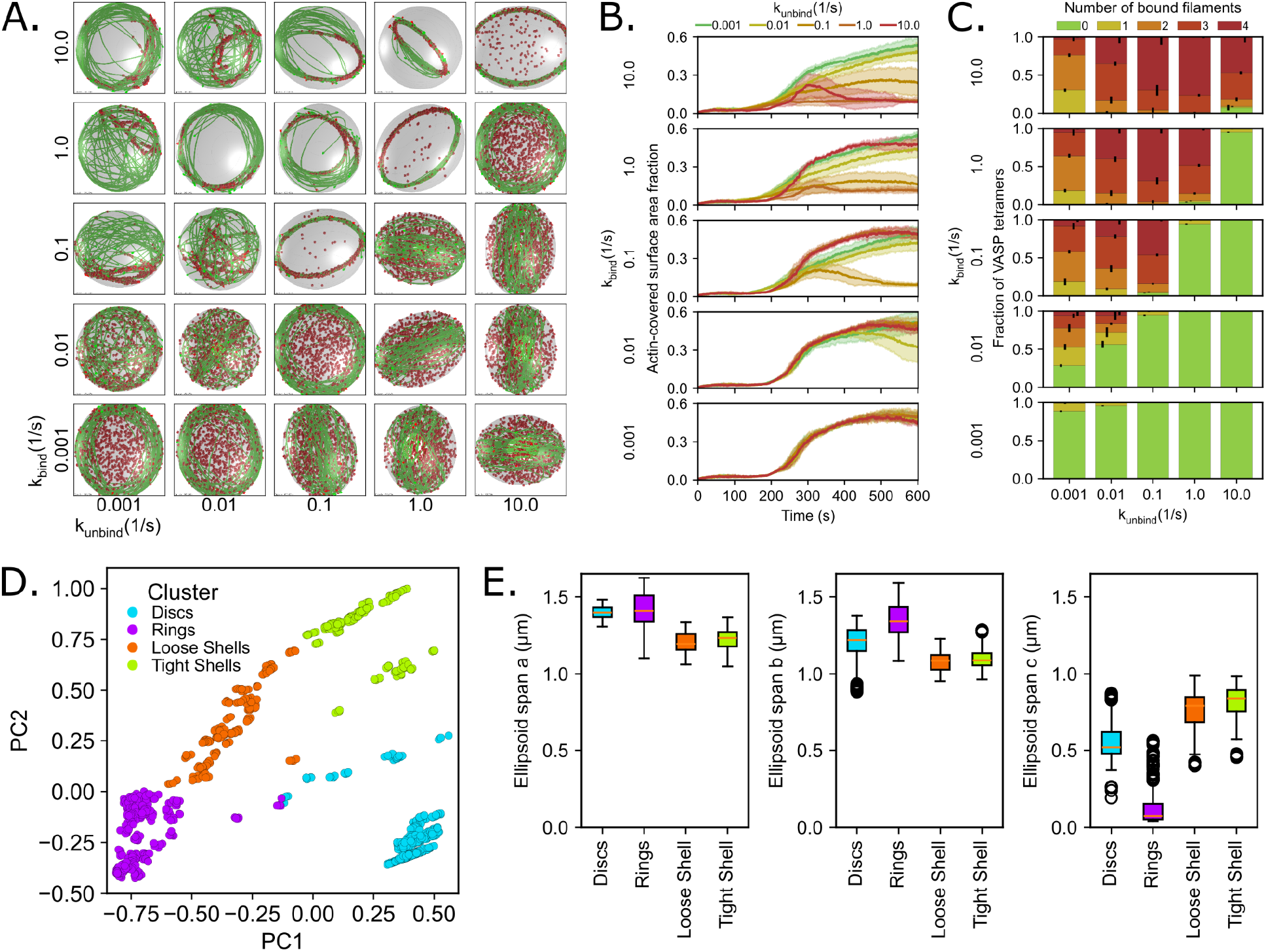
Interactions between VASP binding kinetics and a deformable ellipsoidal droplet reveal the formation of loosely bundled discs. **A)** Representative final snapshots (t = 600 s) from simulations at various binding and unbinding rates within condensates with a deformable ellipsoidal boundary (initially spherical with R = 1 μm) containing 30 actin filaments (green) and 1000 tetravalent crosslinkers (red spheres). The binding rates of the tetravalent crosslinkers are varied along each column, and unbinding rates are varied along each row. The polymerization rate at the plus (+) end is constant at 0.0103 μm/s, and neither end undergoes depolymerization. The deformable boundary has a surface tension of 7 pN/µm and an effective viscosity of 100 pN s/µm. **B)** Time series showing the mean (solid line) and standard deviation (shaded area) of the actin-covered surface area fraction for varied tetravalent crosslinker binding and unbinding kinetics. **C)** Fraction of tetravalent crosslinkers bound to 0, 1, 2, 3, or 4 actin filaments for each condition. The error bars represent the standard deviation. Data was obtained from the last 30 snapshots (5%) of each replicate. **D)** K-means clustering was performed on the last five snapshots from each replicate (data shown in **B** and **C**), which revealed four cluster categories corresponding to actin structures that are discs, rings, loose shells, or tight shells. The data for each snapshot is projected onto a scatterplot of the first two principal components (PC1 and PC2), and colored by clusters. **E)** Ellipsoid span a, span b, and span c (a ≥ b ≥ c) are calculated from the eigenvalues of the gyration tensor and approximate the shape of the actin network as an ellipsoid. The distribution of each ellipsoid span is depicted as box plots and organized by cluster category. Each boxplot displays the median (orange line), interquartile range (box), 95% confidence interval (whiskers), and outliers (open circles).

Finally, we used principal component analysis (PCA) to classify the different actin network structures. PCA showed that for the data in **Figure 2B-C**, which represents five independent dimensions, the first three principal components (PCs) represent ∼96% of the variance in the dataset (**Fig. S2A**). To understand how each of the five parameters used to describe actin organization contribute to each PC, we then looked at the varimax loading parameter. We found that PC1 captured information on the actin-covered surface area fraction and the fraction of VASP tetramers fully bound to four filaments while PC2 captured information regarding the number of VASP tetramers bound to one and two filaments (**Fig. S2B**). Then, using a K-means clustering algorithm on the first three PCs, we classified the resulting actin network shapes into four distinct clusters (**Fig. 2D**). We determined that four is the optimal number of clusters by choosing the largest silhouette coefficient among various cluster sizes (**Fig. S2C**). We computed the eigenvalues of the gyration tensor of the network to extract the ellipsoid spans. This analysis allowed us to approximate the shape of each actin network as an ellipsoid that best described the distribution of filament positions. This analysis showed that the cluster with the lowest span c corresponded to rings, the cluster with an intermediate span c value corresponded to discs, and the two clusters with the highest span c corresponded to both loose and tight shells (**Fig. 2E**). This interpretation matches with a visual inspection, which reveals that these four clusters generally correspond to actin networks that form discs, rings, loose shells, and tight shells. Notably, the cluster corresponding to discs correlated positively with PC1 and negatively with PC2, meaning that the VASP crosslinkers in disc conditions were largely bound to very few to no filaments at all. The absence of a strong contribution from the crosslinkers suggests that disc formation was likely a purely energy minimization phenomenon arising from the interplay between droplet surface energy and filament bending energy rather than the result of VASP crosslinker-mediated kinetic trapping.

### Actin bundle thickness and droplet diameter satisfy a power-law relationship

We next sought to systematically understand the dynamics of droplet deformation. We first analyzed how the aspect ratio of the droplet evolved over time under disc forming conditions (weak VASP binding kinetics, **Fig. 2**, k_bind_ = 0.001 s^-1^) for a droplet of surface tension σ_surface_ = 7 pN/µm. We observed that the aspect ratio evolved non-monotonically with a consistent time signature for all disc-forming conditions. Until the filament length reached the diameter of the droplet, the aspect ratio remained unchanged (L_fil_ = 2R µm, t ≈ 200 s). Once the filaments began growing longer than the droplet diameter, the droplet aspect ratio increased until the filament length reached L_fil_ = πR µm (t ≈ 300 s). As the filaments began to bend around the periphery of the droplet, the aspect ratio decreased, indicating a partial recovery of the spherical shape after an ellipsoidal deformation. However, once the bundles of actin grew longer than πR µm, the aspect ratio returned to growing monotonically (**Fig. 3A, Fig. S3**).

**Figure 3:**
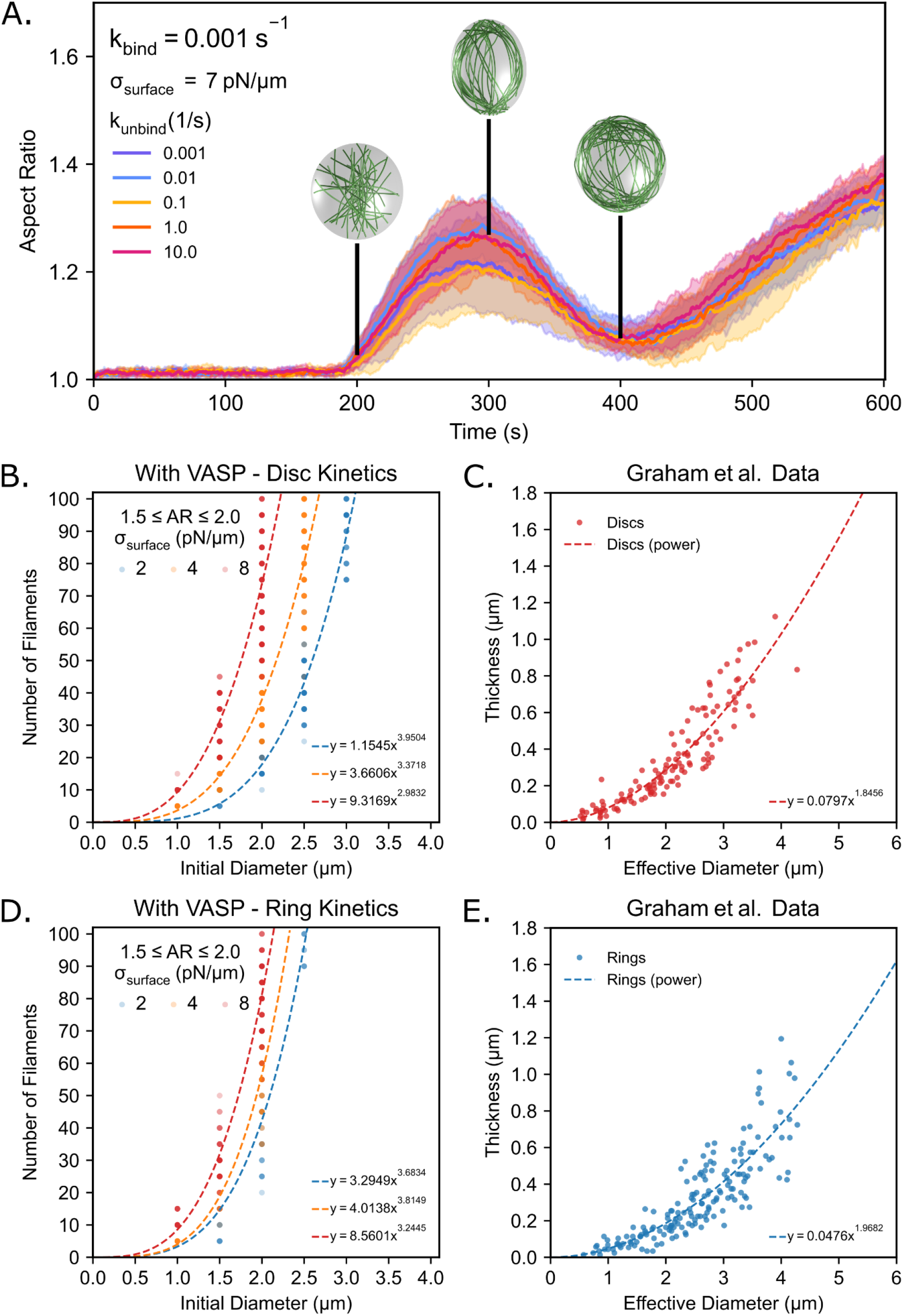
Power law scaling relationship between the initial diameter of the condensate and the number of filaments. **A)** Time series showing the mean (solid line) and standard deviation (shaded area) of condensate aspect ratio for each k_unbind_ condition when k_bind_ = 0.001 s^-1^. The aspect ratio is defined as the ratio between the longest and shortest axes of the ellipsoid (AR = a/c) where a ≥ b ≥ c. The black bars at timestamps t = [200 s, 300 s, 400 s] correspond to when each filament is roughly of length [2π/3 µm, π µm, and 4π/3 µm], respectively. Inset images are visualizations of the filaments and boundary without rendering crosslinkers. **B)** Power law relationship for simulations with VASP that have disc-forming kinetics and with a surface tension of 2 pN/μm (R^2^ = 0.9404, n = 131), 4 pN/μm (R^2^ = 0.8723, n = 179), and 8 pN/μm (R^2^ = 0.7513, n = 170). Data for each power law fit is taken from final condensate aspect ratios at t = 600 s that are between 1.5 and 2.0. **C)** Actin ring thickness as a function of the effective droplet diameter for condensates with aspect ratios < 1.1 (R^2^ = 0.8430, n = 131). Data were taken and reanalyzed from Fig. 3E of Graham et al. 2023^9^. **D)** Power law relationship for simulations with VASP that have ring-forming kinetics and with a surface tension of 2 pN/μm (R^2^ = 0.8190, n = 51), 4 pN/μm (R^2^ = 0.8201, n = 66), and 8 pN/μm (R^2^ = 0.9038, n = 116). Data for each power law fit is taken from final condensate aspect ratios at t = 600 s that are between 1.5 and 2.0. **E)** Actin ring thickness as a function of the effective droplet diameter for condensates with aspect ratios < 1.1 (R^2^ = 0.7278, n = 173). Data were taken and reanalyzed from Fig. 3E of Graham et al. 2023^9^.

From these dynamics, three interesting questions emerge: What is the relationship between droplet size, filament bending energy, and the extent of droplet deformation? What factors control the onset of droplet deformation? And what factors control the dynamics of droplet deformation? We systematically answer these questions below.

To understand the relationship between the droplet size and extent of droplet deformation, we recall that disc formation is governed by an interplay between droplet surface energy and filament bending energy^5,9^. Since the filament bending energy is directly related to the thickness of the filament bundle, varying the number of filaments within our simulations will modulate the accumulation of filament bending energy in the system. Varying the droplet size, on the other hand, will change the droplet surface energy, which is given by the product of the surface tension and the surface area of the droplet. A parameter sweep of the number of filaments versus the initial diameter of the droplet will thus allow us to study their relationship with droplet deformation. To test this idea, we simulated actin bundling in deformable droplets of surface tension σ_surface_ = [2, 4, 8] pN/µm that ranged in size from an initial diameter of 0.5 µm to 4.0 µm and varied the number of filaments from 5 to 100 under fixed VASP tetramer concentration (∼0.40 µM). Additionally, the growth rate of each filament was adjusted so that the final filament length matched the circumference of the original droplet (2πR µm or πD µm) after 600 s for each droplet size. Our choice of surface tension values ensured that we were operating in the deformable regime established in our 30 filament, 2 µm diameter droplet simulations (**Fig. S3**). We simulated 960 unique conditions (10 replicates each, parameters tested shown in **Supplementary Table S2**). To understand how the number of filaments required to produce a small droplet deformation scaled with droplet size, we computed the following. The final aspect ratio was computed for each of the 960 conditions studied and calculated the number of filaments where the droplet aspect ratio fell between 1.5 and 2.0. This range was held constant across all 960 conditions (surface tension values, droplet radii, and number of filaments) studied to provide a common threshold with a substantial sample size.

We first investigated the case of VASP crosslinkers with disc-forming kinetics (k_bind_ = 0.1 s^-1^, k_unbind_ = 1.0 s^-1^). Indeed, we found that the droplet aspect ratio increased with an increase in the number of filaments, and, as the initial diameter of the droplet increased, more filaments were required to deform the droplet to the same or similar aspect ratio. We found that to produce small deformations, the number of filaments and the initial droplet diameter had a power law relationship, of form y = ax^b^, for all surveyed surface tensions for VASP crosslinkers with disc-forming kinetics (**Fig. 3B**). We found that this relationship held even under ring-forming conditions (**Fig. 3D**).

This observation aligns with experimental findings that were shown in our previous work, wherein we found that the thickness of actin rings that deformed to higher aspect ratios increased with increasing droplet diameter^9^. Additionally, VASP droplets that were deformed into higher aspect ratio structures tended to have thicker rings in comparison to actin rings in spherical droplets. In these experiments, a power law relationship was also seen between actin ring thickness and the effective diameter of the VASP droplets. We reanalyzed the experimental data from those prior experiments and restructured it to compare to the model predictions **(Figs. 3C and 3E)**. We found that these experimental results corroborated the computational predictions, and together the model and experimental findings suggest that the thickness of the actin ring follows a power law growth relationship with condensate diameter, and the thickness of the actin ring contributes to the deformation of the VASP droplets. We note that this power law relationship is also consistent with previous theoretical predictions^5^ and energetic arguments^9^. The derivation of this power law relationship can be found in the **Supplementary Materials**. Thus, using our agent-based simulations, we were able to confirm the existence of a power law relationship between the initial diameter of the droplet and the number of actin filaments for different droplet surface tensions.

### Filament length determines the onset and extent of droplet deformation

We next investigated the role of filament length on droplet deformation. In our prior studies, we showed that the filament length controls both the onset of rings and the final aspect ratios as the rings deform the droplet^9,18^. Therefore, we investigated the relationship between filament length and droplet deformation under ring-forming conditions. Further, we studied this relationship in the presence of the actin capping protein, which arrests polymerization of filaments^1^. Specifically, we simulated the stochastic capping and uncapping of actin filaments (**Fig. S4**) using an implicit capping and uncapping framework as follows. In our simulations, plus ends were allowed to switch between capped (at rate k_cap_) and uncapped (at rate k_uncap_) states stochastically throughout the simulation. Once capped, the plus ends could no longer grow until the ends were uncapped. As a result, the filaments reached a distribution of lengths and were crosslinked by VASP tetramers to form rings (k_bind_ = 10.0 s^-1^, k_unbind_ = 1.0 s^-1^) within both rigid (**Fig. S4**) and deformable droplets (**Fig. 4**). In this implicit framework, while we specified the rate of filament (un)capping, we did not specify a total number of capping protein molecules.

**Figure 4:**
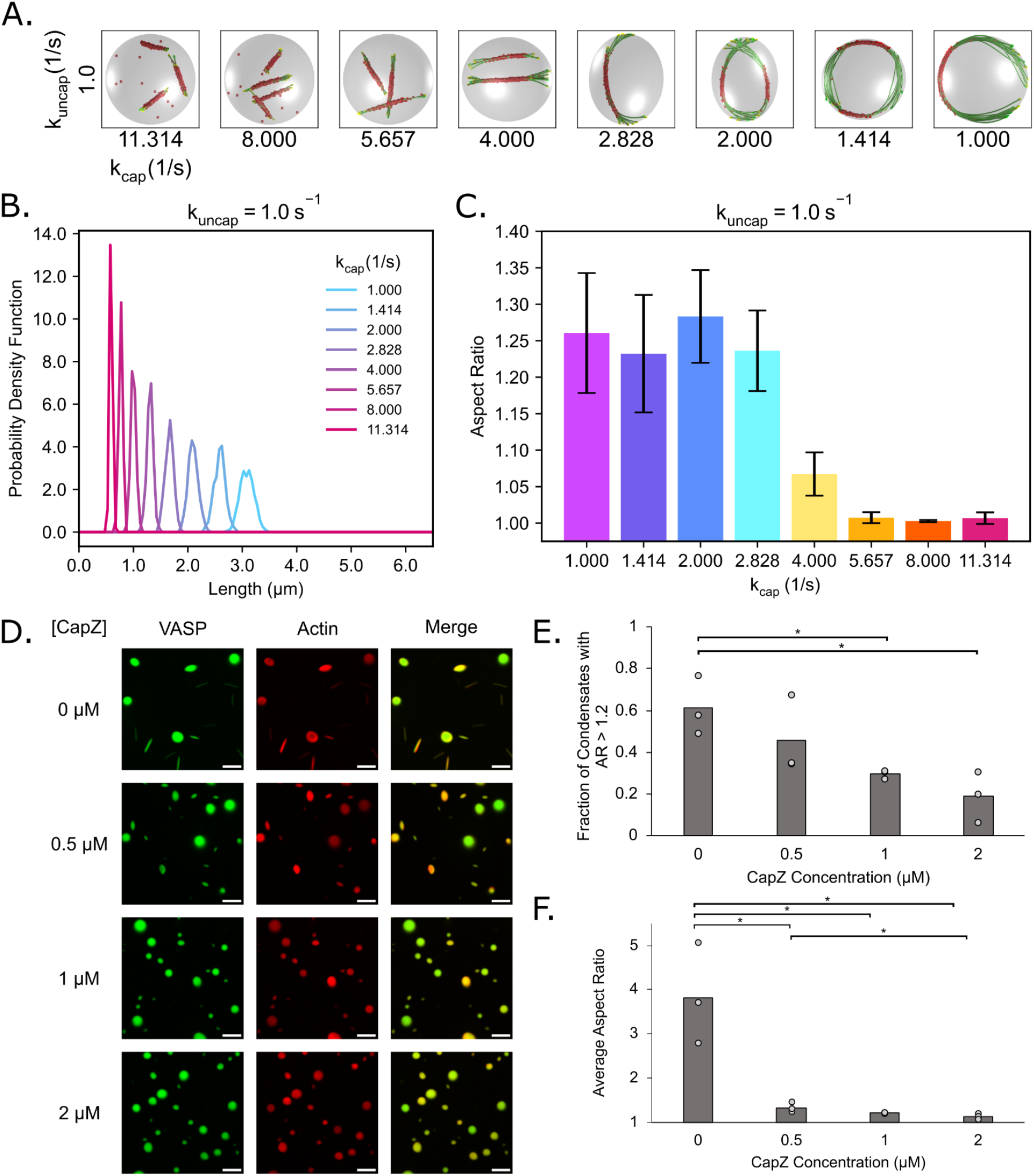
Capping protein tunes filament length and, therefore, the extent of condensate deformation. **A)** Representative final snapshots (t = 600 s) from 10 replicates at various capping and uncapping rates within condensates with a deformable ellipsoidal boundary (initially spherical with R = 1 μm) containing 30 actin filaments (green) and 1000 tetravalent crosslinkers (red spheres). The filament growth rate is fixed at 0.0103 μm/s, and the capping rates are varied while the uncapping rate is held constant at 1.0 s^-1^ to sample the transition between short rods and ring structures. The binding rates of the tetravalent crosslinkers are chosen to promote ring formation when L_fil_ = 2πR µm. The deformable boundary has a surface tension of 4 pN/µm and an effective viscosity of 100 pN s/µm. **B)** Probability density function of the filament length for each simulation condition associated with **A**. As the capping rate increases, the probability of finding longer filaments decreases. **C)** Bar chart showing the mean final condensate aspect ratio for each condition. The aspect ratio is defined as the ratio between the longest and shortest axes of the ellipsoid (AR = a/c), where a ≥ b ≥ c. **D)** Fluorescence images of condensates (3 µM Atto 594 labeled actin and 10µM Atto 488 labeled VASP) with increasing CapZ concentrations [0 µM, 0.5 µM, 1 µM, and 2 µM] showing the suppression of high aspect ratio condensates as CapZ concentration increases. Scale bars, 5 µm. **E)** Quantification of the fraction of condensates with an aspect ratio greater than 1.2 for the conditions shown in **D**. Data are the mean across three independent experiments. The overlaid gray circles denote the means of each replicate. One asterisk denotes p <.05 using an unpaired, two-tailed t-test on the means of the replicates, n=3. **F)** Quantification of the average aspect ratio for conditions shown in **D**, showing a decrease in aspect ratio as CapZ concentration increases. Data are the mean across three independent experiments. The overlaid gray circles denote the means of each replicate. One asterisk denotes p <.05 using an unpaired, two-tailed t-test on the means of the replicates, n=3. Buffer conditions for all conditions were 20 mM Tris pH 7.4, 150 mM NaCl, 5 mM TCEP, and 3% (w/v) PEG 8000.

We varied the capping rates in the range [k_cap_ = 1.0 s^-1^ - 11.314 s^-1^] for fixed uncapping rate k_uncap_ = 1.0 s^-1^ (**Fig. 4A**). These values were chosen because they produced the full range of VASP-crosslinked actin structures from linear bundles to rings (**Fig. S4**). We found that increasing capping rate (k_cap_) resulted in shorter filaments and a narrower distribution of filament lengths (**Fig. 4B**). Additionally, increasing k_cap_ also resulted in a narrower filament length distribution but centered around a smaller filament length (**Fig. 4B** and **Fig. S4D**). We quantified the aspect ratio of the droplet as a measure of the extent of the deformation. Indeed, increasing the capping kinetics resulted in lower aspect ratio droplets because shorter filaments cannot exert large deforming forces on the droplet surface (**Fig. 4C**). From these simulations, we predicted that when capping protein was added to VASP droplets with actin in them, the average droplet aspect ratio would decrease and there would be fewer droplets with high aspect ratio.

To test this prediction in experiments, we incorporated capping protein, CapZ, into droplets composed of VASP. CapZ is an actin-binding protein that regulates filament dynamics by binding tightly to the barbed ends of actin filaments, thereby preventing further monomer addition and elongation and effectively controlling filament length ^30–32^. Specifically, we formed droplets in a 10 μM solution of VASP in the presence of 3 μM Actin with increasing concentrations of CapZ. Droplets were formed by mixing VASP and associated proteins with 3% (w/v) PEG 8000 as a crowding agent. PEG is commonly added in the study of LLPS to mimic the crowded environment in the cell cytoplasm ^33–35^. As was observed previously^9,17,18^, VASP droplets lacking CapZ deformed into rod-like shapes upon actin polymerization (**Fig. 4D**). As increasing concentrations of CapZ were added, the droplets became more spherical (**Fig. 4D**), and both the fraction of deformed condensates, defined as protein droplets with an aspect ratio greater than 1.2, and the average aspect ratio of droplets decreased with increasing CapZ concentration. (**Fig. 4E-F**). Notably, phalloidin-stained actin did not arrange into rings at the inner surfaces of droplets in the presence of CapZ (**Fig. S4E**), suggesting that CapZ addition reduced actin filament length such that there were no filaments long enough to exceed the condensate diameter.

### Dynamics of droplet deformation shows a dynamic snapping behavior depending on the droplet surface tension

We previously showed that the dynamics of droplet deformation for deformable VASP droplets can be non-monotonic in time (**Fig. 3A**). To investigate how these dynamics depend on VASP and droplet properties, we tracked the droplet deformation dynamics for different VASP binding-unbinding rates and for different values of droplet surface tension (σ_surface_ = 7 pN/µm (**Fig. 5A-C**), and σ_surface_ = 2 pN/µm (**Fig. 5D-F**)). Droplet deformation can be understood as the balance between the bending energy of the actin bundle and the surface energy of the droplet. The onset of droplet deformation occurs when the length of the bundle exceeds the diameter of the droplet because the linear actin bundles are able to overcome the surface energy of the droplet. However, the extent of droplet deformation depends on VASP-actin interaction kinetics. When k_bind_ > k_unbind_, actin in ring-like states with strongly-bound VASP tetramers leads to minimal droplet deformation. These rings are kinetically trapped, which results in reduced deformation capacity of the actin networks (**Fig. 5B-C, Movie M1**).

**Figure 5:**
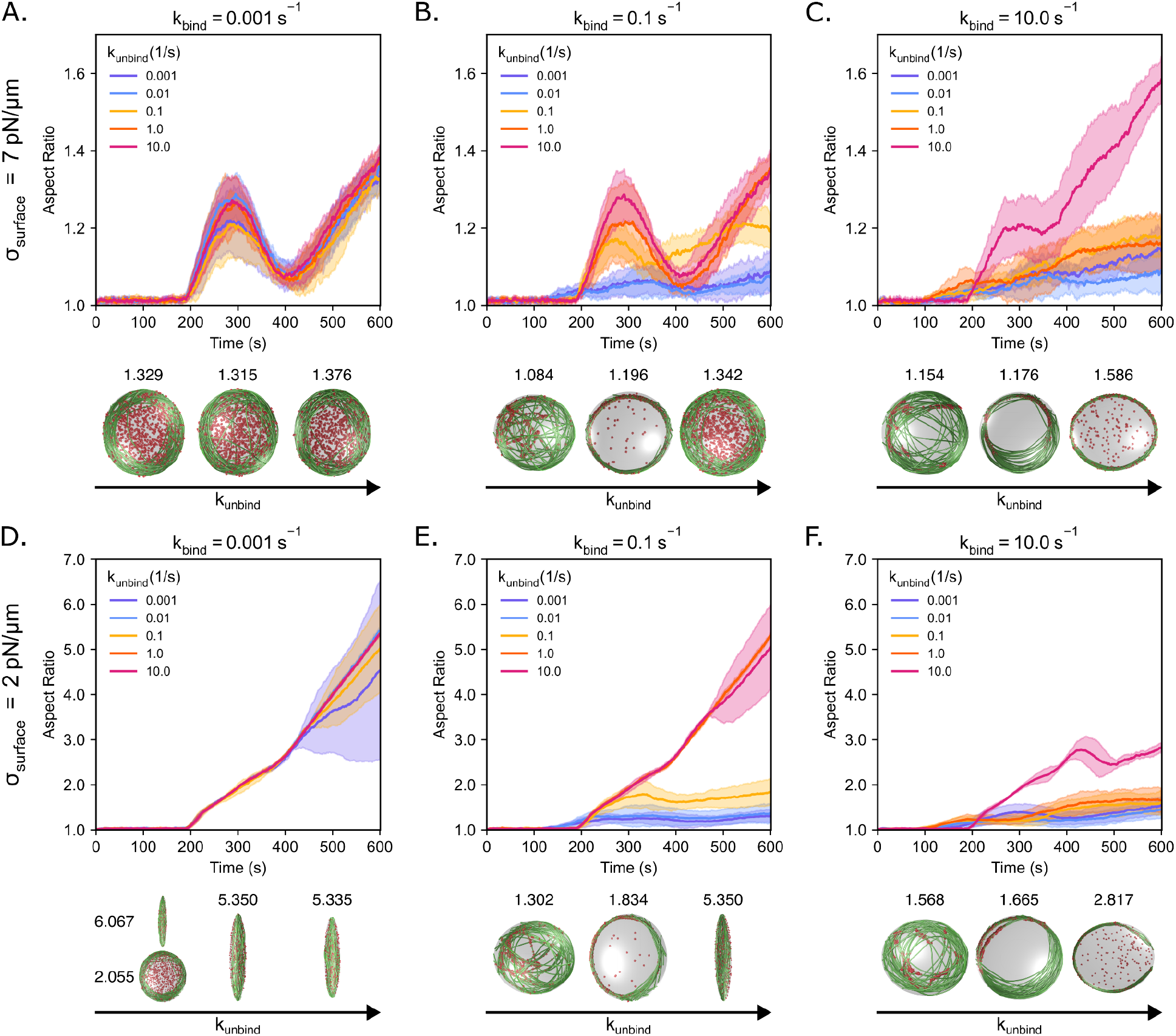
VASP crosslinker binding and unbinding kinetics alter the dynamics of droplet deformation governed by the competition between filament bending energy and condensate surface energy. Time series showing the mean (solid line) and standard deviation (shaded area) of condensate aspect ratio for each condition. The aspect ratio is defined as the ratio between the longest and shortest axes of the ellipsoid (AR = a/c), where a ≥ b ≥ c. For surface tension σ_surface_ = 7 pN/µm: **A)** k_bind_ = 0.001 s^-1^, **B)** k_bind_ = 0.1 s^-1^, and **C)** k_bind_ = 10.0 s^-1^. For surface tension σ_surface_ = 2 pN/µm: **D)** k_bind_ = 0.001 s^-1^, **E)** k_bind_ = 0.1 s^-1^, and **F)** k_bind_ = 10.0 s^-1^. Please note that the aspect ratio scales are different between panels A-C and D-F. The snapshots show actin in green and VASP in red. High aspect ratio droplets have almost linear rod-like actin bundles. The snapshots within each panel correspond to k_unbind_ values 0.001 s^-1^, 0.1 s^-1^, and 10.0 s^-1^. Final aspect ratios are mentioned above each of the snapshots.

When k_bind_ ≤ k_unbind_, we observed an increase in droplet deformation depending on the value of the surface tension. Under disc-forming conditions, where actin bundling is weak, the droplet begins to deform once the filaments reach L_fil_ > 2R (t ∼ 200 s). As the filaments grow longer, the filaments begin deforming the droplet linearly with time for low surface tension (σ_surface_ = 2 pN/µm) to form rod-like bundles of actin (**Fig. 5D-E, Movie M2**). However, the variability increases with time in low surface tension and low binding and unbinding rates. In this case, we observed that out of ten replicates, six of them were rods while four of them snapped back into discs (**Fig. 5D**, left-side snapshots). On the other hand, when σ_surface_ = 7 pN/µm, the droplet aspect ratios evolve as a non-monotonic hump (t ∼ 200-400s), after which they either experience continuous linear growth, or remain stable. This nonlinear temporal behavior can be understood as follows: the aspect ratio of the droplet increases initially with filament growth as filaments are aligned along ellipsoidal major axis (**Movie M3**). This anisotropic filament distribution enables for filament growth-driven droplet deformation. As the filament length reaches L_fil_ = πR µm at t∼300s, the filaments bend along the periphery of the droplet, resulting in spindle shaped networks that trace the contour of the ellipsoidal droplet forming an near-isotropic network (**Movie M3**). In this state, actin does not drive droplet deformation as the filament segments are arranged isotropically along the droplet surface. The actin filaments align themselves and grow along the droplet surface droplet surface; such tangential growth does not lead to any forces exerted by the actin bundle on the droplet interface. As a result, the droplet surface energy overcomes bending energy and the droplet tends towards a spherical state. Eventually (t > 400 s, L_fil_ = 4πR/3 µm), the filaments rearrange into discs that can overcome the surface energy resulting in droplet deformation towards an ellipsoidal shape. This can be observed through the reduction in actin-covered surface area in t ∈[300,400] s as shown in **Fig. 2B**. From ∼200-400 s, the initial increase and decrease in aspect ratio forms a hump in the graph that can be understood in the context of a dynamic snapping behavior. Snap-through bifurcations are a common feature in mechanics when small changes to a control parameter can lead the system from one stable energy minimizing state to another^36–38^. However, in our case, we observe a dynamic snapping because the kinetic rearrangement of the actin filaments over time leads to a time-dependent response.

In the case of ring formation kinetics, the stronger actin binding-unbinding kinetics, specifically when k_bind_ ≈ k_unbind_ for k_bind_ ≥ 0.1 (**Fig. 5B-C**), result in actin bundles that have formed by the time that the filament length reaches the diameter of the droplet, L_fil_ = 2R µm (t ≈ 200 s). By forming a bundle early, nascent ring structures emerge before L_fil_ = πR µm (t ≈ 300 s) in the form of either a single main bundle or two primary bundles oriented 180° from each other. Here, the filaments are arranged into the bundle to minimize bending energy and, with the guidance of crosslinkers that favor bringing filaments together, are better able to concentrate force to deform the droplet. However, crucially, because the filaments have generally been confined into a highly-aligned, essentially 2D ring structure stabilized by crosslinkers, the bending energy of the ring is able to overcome the droplet surface energy to push the droplet toward higher aspect ratios without experiencing a hump as was the case for discs.

### Deformable droplets made of ABPs that form dynamic multimers also exhibit actin bundling and dynamic snapping

We next investigated whether the relationship between droplet deformation and actin bundling was specific to VASP alone or generalizable to other ABPs. To do so, we investigated the role of monomeric crosslinkers that can associate and dissociate dynamically with other monomers to form dynamic multimers in deforming the droplet. We simulated ABPs as monomeric units capable of binding to a single actin filament and up to two other monomers, thus allowing for the formation of higher-order multimers (**Fig. 1G**)^8^. We prescribed ring-forming actin-binding kinetics while modulating the kinetics of multimer formation and splitting and found that, actin rings and discs can form depending on the different kinetic parameters (**Fig. 6A, Fig. S5**). Our analysis of multimer lengths showed that rings only formed in kinetic conditions that favored multimerization of the crosslinkers, while discs formed under conditions in which the crosslinkers remained largely monomeric (**Fig. S5D**).

**Figure 6:**
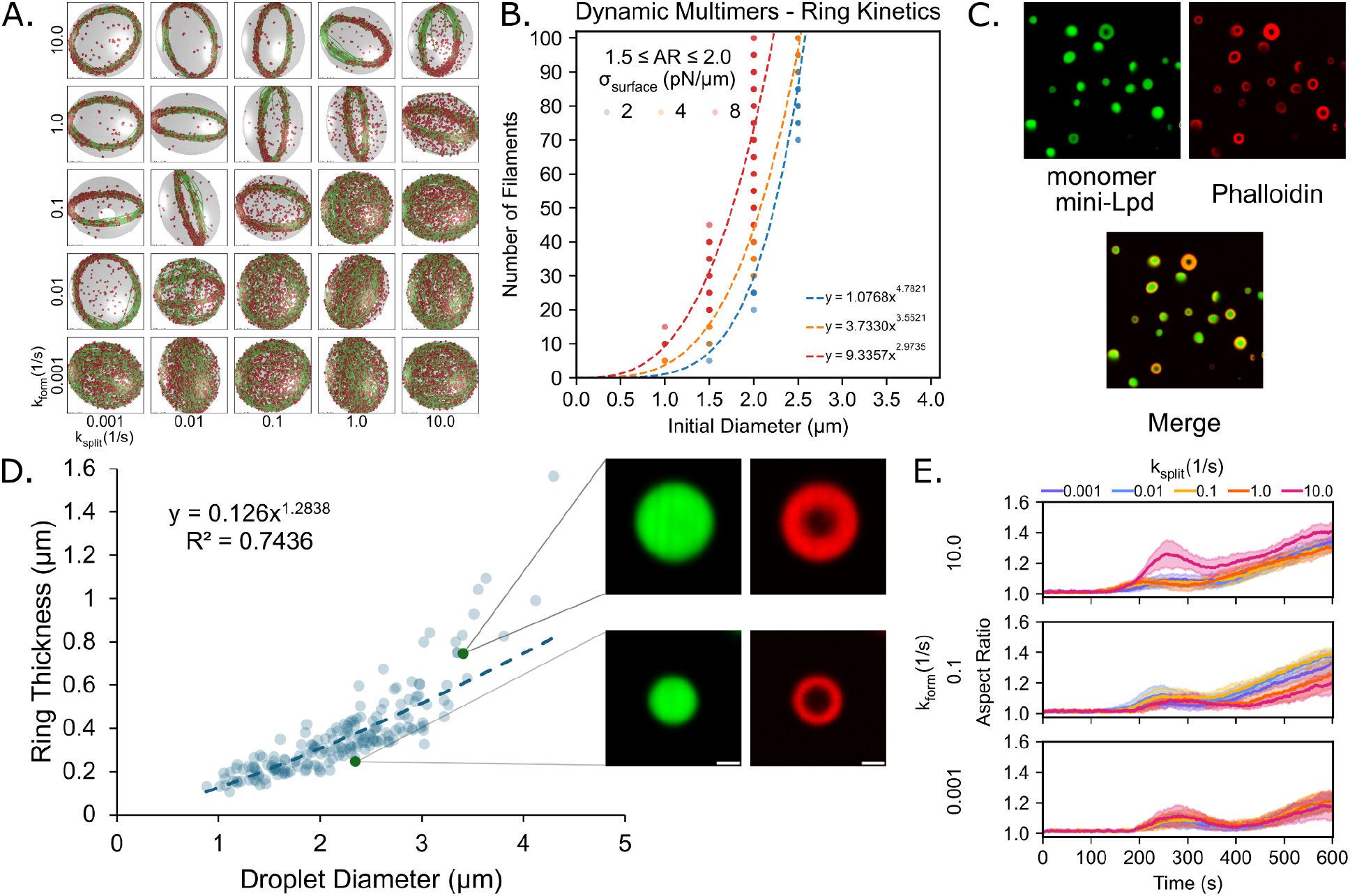
Multivalent condensate environments exhibit disc formation and a power law scaling relationship between the initial diameter of the condensate and the number of filaments. **A)** Representative final snapshots (t = 600 s) from simulations at various multimer forming and splitting rates within condensates with a deformable ellipsoidal boundary (initially spherical with R = 1 μm) containing 30 actin filaments (green) and 2000 monomeric crosslinkers (red spheres) which can dynamically form multimers of various lengths. The monomer-monomer binding rates of the tetravalent crosslinkers are varied along each column, and monomer-monomer unbinding rates are varied along each row. The actin-binding rates of the monomers are consistent with ring-forming kinetics determined from previous simulations. The polymerization rate at the plus (+) end is constant at 0.0103 μm/s, and neither end undergoes depolymerization. The deformable boundary has a surface tension of 7 pN/µm and an effective viscosity of 100 pN s/µm. **B)** Power law scaling relationship between the number of filaments and the initial diameter of deformable condensates in dynamic multimerization simulations with a surface tension of 2 pN/μm (R^2^ = 0.9377, n = 103), 4 pN/μm (R^2^ = 0.9440, n = 104), and 8 pN/μm (R^2^ = 0.7472, n = 173). **C)** Representative confocal cross-section images of condensates formed from 5 µM monomer-mini-Lpd after the addition of 3 µM actin and staining with Phalloidin iFluor 594 showing rings of polymerized actin. Buffer conditions were 20 mM Tris pH 7.4, 50 mM NaCl, 5 mM TCEP, and 3% (w/v) PEG 8000. Scale bars, 5 µm. **D)** Power law scaling relation between the measured actin ring thickness and the diameter of monomer mini-Lpd condensates. The inset images correspond to the denoted points on the graph and depict examples of the difference in measured actin ring thickness. Scale bars, 1 µm. **E)** Time series showing the mean (solid line) and standard deviation (shaded area) of condensate aspect ratio for each condition. The aspect ratio is defined as the ratio between the longest and shortest axes of the ellipsoid (AR = a/c), where a ≥ b ≥ c.

We also validated the presence of power law relationship between droplet size and number of filaments. We chose the multimerization condition that maximizes crosslinker multimer size (**Fig. S5D**, k_form_ = 10.0 s^-1^, k_split_ = 0.01 s^-1^). We found that the initial radius of the droplet and the number of filaments required to deform it also followed a power law relationship (**Fig. 6B**) similar to stable tetramers studied earlier (**Fig. 3B, 3D**). To test our model prediction that dynamic multimers of ABPs were sufficient to bundle actin filaments and sustain droplet deformation within protein condensates, experiments were performed using condensates made of an EGFP-tagged c-terminal IDR region of the Lamellipodin protein. This protein, henceforth called monomer-mini Lpd as it is a native monomer, was shown to phase separate and facilitate actin filament assembly^8^. To test if there was a power law relationship between actin ring thickness and condensate diameter, we performed similar experiments to those done previously with VASP (**Fig. 3C, 3E**)^9^ with condensates composed of monomer mini-Lpd. Specifically, condensates were formed from a 5 µM solution of monomer mini-Lpd and 3 w/v% PEG 8000. Monomeric actin (G-actin) was added to the condensates at a concentration of 3 µM and allowed to assemble prior to filamentous actin staining with phalloidin-594 (**Fig. 6C**). Actin rings appeared at the droplet periphery. Actin ring thickness within mini-Lpd condensates was quantified by Gaussian fitting of the fluorescence intensity profile across one side of each ring to measure the full-width at half-maximum, subtracting the diffraction limit to approximate the true thickness of the actin rings. The effective spherical condensate diameter for asymmetrical condensates was calculated by plotting ring thickness as a function of condensate diameter (**Fig. 6D**). Ring thicknesses somewhat below the optical diffraction limit were estimated by measuring the Gaussian intensity profile for actin at the droplet edge and subtracting the microscope point spread function. In line with model predictions, condensates formed from monomer mini-Lpd also displayed a power law relationship between condensate diameter and actin ring thickness, displaying similar values in the power law fit to those displayed in VASP condensates. Together, the experimental results and modeling predictions suggest that this power law relationship may hold across any condensate-forming protein that facilitates actin filament assembly.

For simulations, analyzing the deformation kinetics of these droplets showed that dynamic multimer crosslinkers promoted a higher final aspect ratio than the analogous condition for stable tetramer (VASP) (compare **Fig. 6E** and **Fig. 5C**). Furthermore, while all conditions experienced droplet deformation, the conditions that formed rings had larger final aspect ratios than the conditions that formed discs (**Fig. 6E, Fig. S5C**). Thus, dynamic multimers form robust rings with additional modes of reorganization, which facilitates the bending energy dissipation resulting in higher aspect ratios. This also alters the dynamics of droplet deformation as reflected by the lack of pronounced hump in aspect ratio time series (**Fig. 6E, Fig. S5C**).

### Confinement of growing actin filaments in liquid droplets is sufficient for filament bundling and droplet deformation in the absence of crosslinkers

Our analysis of VASP and Lpd droplets and their interactions with actin shows that loosely bundled actin discs emerge when the droplet boundary is allowed to deform, even when crosslinker interactions with actin filaments are minimal. The lack of appreciable crosslinker activity suggests that disc formation might be primarily driven by the material properties of the droplet itself, rather than the crosslinkers. Therefore, we next investigated how droplet properties impacted actin bundling. Specifically, we varied σ_surface_ and µ_effective_ in a deformable droplet in the absence of any crosslinking interactions. We found that actin filaments formed loose discs when the condensate surface tension was sufficiently low (**Fig. 7A**). The fraction of the condensate surface covered by actin was larger for shells than for discs, and the transition from discs to shells was clear as surface tension increased (**Fig. S6A**). We also observed that shell structures were recovered as surface tension increased, indicating that condensate deformation could drive assembly of loosely bundled actin discs. As expected, changes in the effective viscosity had a very muted effect on the final aspect ratio, as this parameter alters only the time necessary to deform the droplet and not the final state of deformation within the deformable ellipsoid framework^26^. Thus, we focused on σ_surface_ for the next steps.

**Figure 7:**
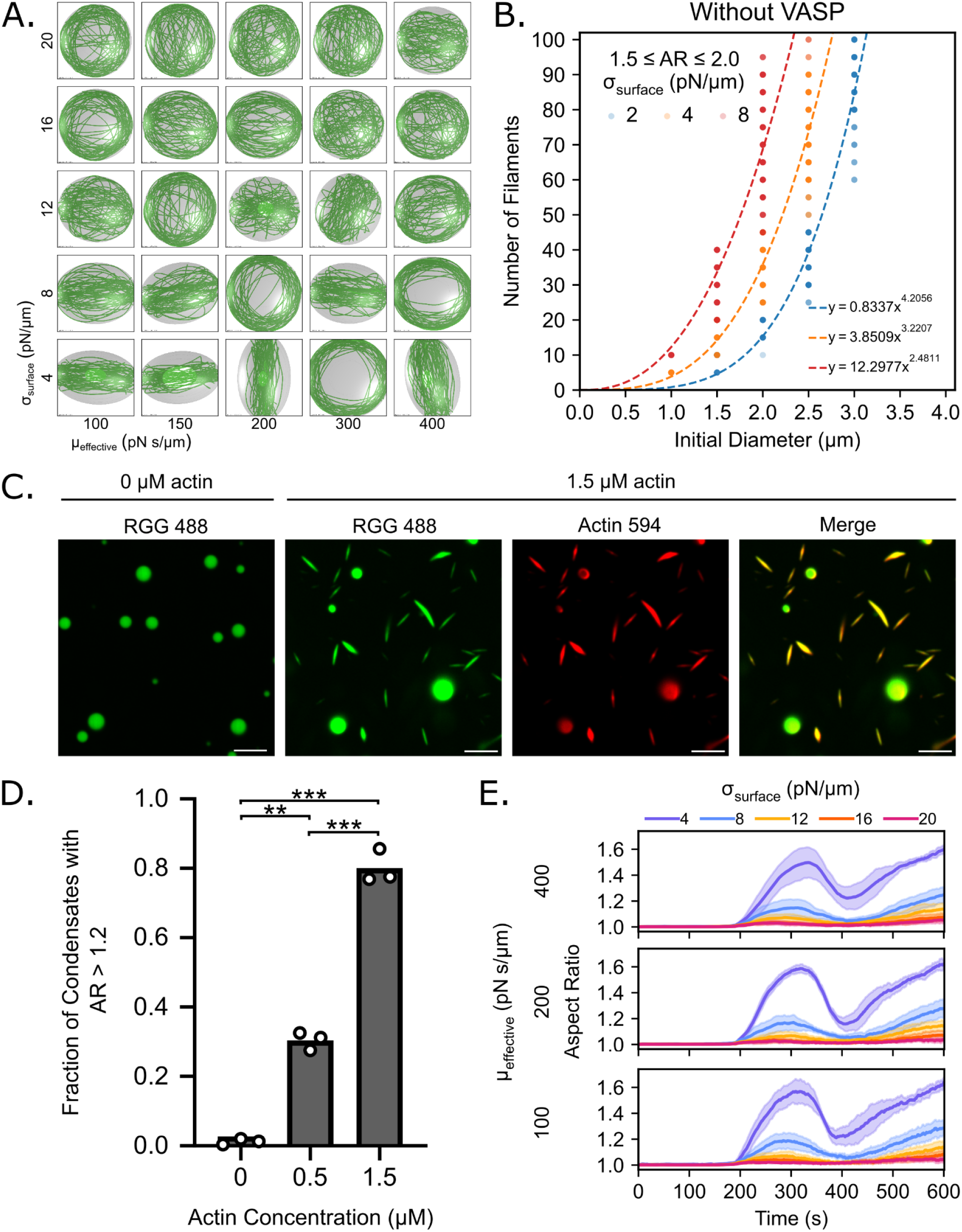
Interfacial properties of the condensate surface govern deformation and determine whether the actin network will form shells or discs. **A)** Representative final snapshots (t = 600 s) from simulations varying the properties of the deformable condensate boundary (initially spherical with R = 1 μm) containing 30 actin filaments (green) and in the absence of any actin crosslinker. The effective viscosity which attenuates the rate of deformation is varied along each column, and the surface tension which describes the innate resistance of the condensate boundary to deformation is varied along each row. The polymerization rate at the plus (+) end is constant at 0.0103 μm/s, and neither end undergoes depolymerization. **B)** Power law relationship for simulations without VASP present and with a surface tension of 2 pN/μm (R^2^ = 0.9230, n = 146), 4 pN/μm (R^2^ = 0.8790, n = 165), and 8 pN/μm (R^2^ = 0.7563, n = 158). Data for each power law fit is taken from final condensate aspect ratios at t = 600 s that are between 1.5 and 2.0. **C)** Fluorescence images of condensates (20µM Atto 488 labeled RGG) with 0 µM and 1.5 µM Atto 594 labeled actin, showing the lack of deformation in the absence of actin and high aspect ratio condensates in the presence of actin. Scale bars, 5 µm. **D)** Quantification of the fraction of condensates with an aspect ratio greater than 1.2 for the conditions shown in **C**, showing an increase in aspect ratio fraction with increasing actin concentration. Data are the mean across three independent experiments. The overlaid white circles denote the means of each replicate. Two asterisks denote p <.01 and three asterisks denote p <.001 using an unpaired, two-tailed t-test on the means of the replicates, n=3. Buffer conditions for all conditions were 20 mM Tris pH 7.4, 50 mM NaCl, 5 mM TCEP, and 3% (w/v) PEG 8000. **E)** Time series showing the mean (solid line) and standard deviation (shaded area) of condensate aspect ratio for each condition. Surface tension is varied within each plot with the same effective viscosity. The aspect ratio is defined as the ratio between the longest and shortest axis of the ellipsoid (AR = a/c) where a ≥ b ≥ c.

We next investigated whether droplet deformation in the absence of crosslinkers would also exhibit a power law relationship between the number of actin filaments and condensate diameter. As before, we varied the number of actin filaments and the initial diameter of the droplet and were able to extract a power law relationship for nonspecific crosslinkers (**Fig. 7B**). Again, we observed that actin elongation alone was sufficient to deform the droplet. This result led us to predict that an arbitrary condensate-forming protein with a weak affinity for actin filaments would be sufficient to form these discs. To experimentally test this prediction, we formed condensates from the RGG domain of LAF-1, a well-characterized phase-separating protein that lacks actin-binding domains^39^, which has a high concentration of positively charged arginine residues and is therefore likely to interact nonspecifically with negatively charged actin. We examined how the addition of actin impacted RGG condensate shape and filament organization. RGG droplets were formed from a 20 µM solution of the protein and 3 w/v% PEG 8000. Increasing concentrations of actin were added to the droplets after they formed, as described above. As predicted by the simulations, RGG condensates were progressively deformed into rod-like morphologies as the concentration of actin increased (**Fig. 7C**). Additionally, both the fraction of deformed condensates, defined as protein droplets with an aspect ratio greater than 1.2 (**Fig. 7D**), and the average aspect ratio of droplets increased with increasing actin concentration. (**Fig. S6C**). Notably, phalloidin-staining revealed that actin filaments were arranged into rods and peripheral rings at the inner surfaces of the droplets (**Fig. S6D**). These experimental observations suggested that actin filament assembly within condensates was alone sufficient to promote the bundling of actin filaments into discs, and that asymmetries in the condensate boundary could alone facilitate bundling of actin filaments into discs. Finally, when we increased the surface tension in the simulations, we observed that the droplets deformed to lower aspect ratios. The dynamics of droplet deformation transitioned from dynamic snapping to slow monotonic increase over time (**Fig. 7E, Fig. S6B**).

### Feedback between droplet shape and bundle alignment is tuned by crosslinker properties

Throughout our simulations, we found different dynamics for droplet deformation depending on the crosslinker kinetics and multivalency and droplet surface tension (**Fig. 3A, 4C, 5, 6E, 7E, S3C, S5C, S6B**). Based on these observations, we hypothesized that there exists a feedback between the droplet shape and filament alignment. To test this hypothesis, we first simulated actin filament elongation and bundling within ellipsoidal rigid droplets without any crosslinkers (**Fig. 8A**). We defined an alignment angle of the actin bundle as the angle formed between ellipsoid span a and span c, obtained from the eigenvalues of the gyration tensor as in **Figure 2E** and described in the Supplementary Methods. A filament alignment angle approaching 0 degrees describes a flat distribution of actin as expected for a ring, while an alignment angle approaching 45 degrees describes a perfectly spherical distribution of actin as expected for a spherical shell. We found a strong relationship between the droplet aspect ratio and filament angle (**Fig. 8**). To understand the role of boundary shape and actin organization, we simulated actin networks within droplets of various aspect ratios (**Fig. 8A**). We found that the actin filaments aligned with the major axis of the ellipsoid at all aspect ratios studied (**Fig. 8A**). When we repeated these calculations with deformable droplets without crosslinkers, we found that decreasing surface tension resulted in both higher aspect ratio droplets and lower filament alignment value (**Fig. 8B**). Further, we observed that small changes to alignment angle resulted in large changes to aspect ratios **(Fig. 8B, inset)**. This observation further validates our hypothesis that the filament alignment angle controls the extent of droplet deformation. When we analyzed the relationship between filament alignment angle and aspect ratio in the presence of dynamic multimers under ring-forming conditions, we saw a shift towards more aligned filaments and higher aspect ratios as multimer formation became more favorable (**Fig. 8C**). When we used stable tetrameric VASP as a crosslinker, we saw that, depending on the binding kinetics, we could achieve the same alignment angle for a wider range of aspect ratios (**Fig. 8D**). Comparing time profiles of aspect ratio and alignment angle evolution between ring forming dynamic multimeric (**Fig. 8C**) and stable, tetrameric (**Fig. 8D**) crosslinkers, we observed that the stability of a multimeric unit also tuned the maximum aspect ratio achieved during deformation. Thus, we concluded that, while the material properties of the droplet determine the extent of filament alignment and droplet deformation, the biochemical properties of ABPs controlled the dynamic emergence of different actin networks.

**Figure 8:**
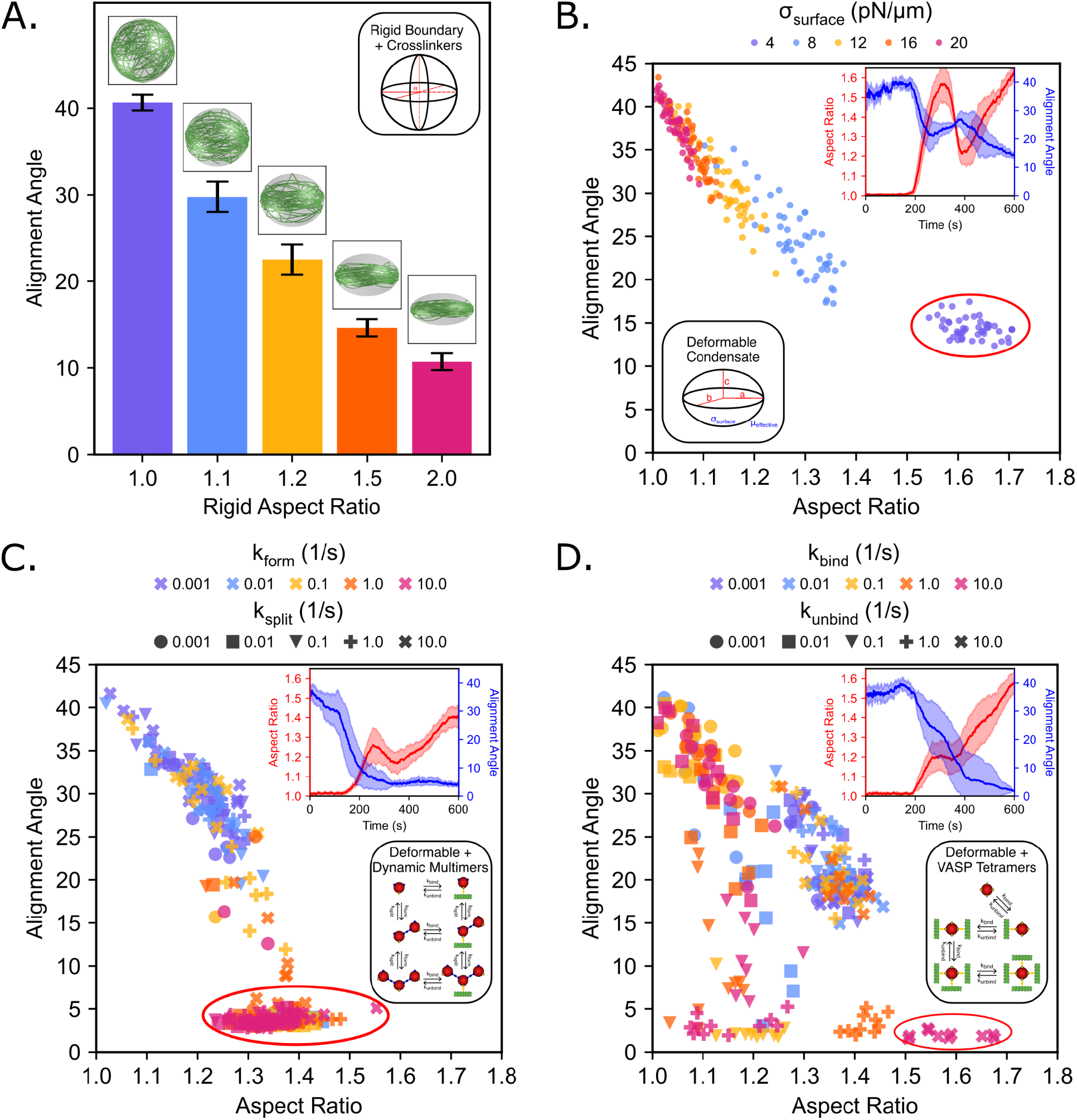
Feedback between droplet deformation and actin filament alignment. **A)** Alignment angle for rigid ellipsoidal droplets with fixed but varied aspect ratios. Inset images are final representative snapshots of each condition. **B-D)** Phase diagrams plotting the final alignment angle and final aspect ratio for each condition across all replicates for simulations: **B)** without crosslinkers (σ_surface_ indicated by color), **C)** with dynamically multimerizing crosslinkers (k_form_ indicated by color and k_split_ indicated by marker shape), and **D)** with VASP crosslinkers (k_bind_ indicated by color and k_unbind_ indicated by marker shape). Inset graphs show the time profile of the alignment angle (blue) and the aspect ratio (red) for the conditions indicated by a red circle: **B)** σ_surface_ = 4 pN/µm, µ_effective_ = 100 pN s/µm, **C)** k_form_ = 10.0 s^-1^, k_split_ = 10.0 s^-1^, **D)** k_bind_ = 10.0 s^-1^, k_unbind_ = 10.0 s^-1^.

## Discussion

In this work, we used a combination of agent-based modeling and in vitro biophysical experiments to systematically dissect how the biochemical and physical properties of liquid-like droplets of ABPs modulate actin filament organization and droplet shape. We show that deformable boundaries can bundle actin through confinement to form either tight rings or weakly-bundled discs characterized by spatially restricted actin filaments. Consequently, the bending energy of the disc or the ring deforms droplets to form ellipsoids. The extent of the droplet deformation depends on both the interfacial properties of the droplet and on the actin network. Filaments in tightly bundled actin rings are unable to unravel to generate large forces on the droplet interface, Thus, kinetically trapped actin rings can only result in moderate droplet deformation. Additionally, multivalent crosslinkers can allow for reorganization leading to higher aspect ratio droplets. On the other hand, actin filaments in weakly-bundled discs can unravel into linear filaments thereby maximizing droplet deformation to form rods. In both cases, the number of filaments needed to deform droplets scales as a power law with respect to droplet radius. These findings provide a mechanistic view of how filament growth, bundling, and droplet deformability give rise to emergent properties of actin networks in liquid-like droplets and a holistic framework to understand previous experimental observations^8,9,12^. Thus, our work establishes the mechanochemical principles of droplet deformation as a function of the properties of droplets and actin networks.

Our finding that confinement alone is a bonafide mechanism of both filament bundling and droplet deformation has implications for how phase separation may organize the cytoskeleton. Previous studies have shown that actin confinement within rigid spheres^17,18^ and inter-filament attraction as necessary conditions for bundling^4,6^. Other studies have shown that confinement of actin in lipid bilayer vesicles can also lead to filament organization^6,40,41^. Here, we highlight that even liquid deformable boundaries offer mechanical feedback to reorganize weakly-bundled filaments into discs. Additionally, consistent with results shown in^4^, we see that rings and discs organize to minimize bending energy within spheroidal droplets. More importantly, the feedback mechanism between droplet deformation and alignment of the actin bundle has implications for directional force generation of actin filaments in cells^42–45^.

Interestingly, when we introduce specific crosslinkers with known actin binding properties such as mini-Lpd or VASP, we expand the parameter space for the formation of actin rings and discs. It is possible that specific ABPs based on their kinetic interaction with actin can lead to a wide range of bundling morphologies with distinct force-generation capacity. Such crosslinking dynamics also have implications for the mechanical properties of the actin networks^1,46,47^.

Looking across all the combinations of cases we have investigated, we found that the relationship between the kinetics of droplet deformation has a rich array of behaviors that can be classified as: no change in aspect ratio, monotonic increase in aspect ratio, and intriguingly, a nonmonotonic dynamic behavior. The common principle among all the droplet deformation dynamics is that the droplet deforms when the energy exerted by the filament bundle overcomes the interfacial energy of the droplet. This energy balance is consistent with what has been observed in other systems of deformable droplets^26^. Furthermore, the dynamics of droplet deformation mimics a dynamic snap-through, where at each time point, the balance of forces dictates the droplet shape but the viscosity of the droplet sets a timescale of deformation^37,38^. Dynamic snap-through is observed in viscoelastic materials or when inertia is added to quasi-static equations of motion in mechanics^37,38^. In our case, we find that the kinetics of actin and crosslinker binding and unbinding adds additional degrees of freedom, leading to the emergence of multiple endpoint morphologies. However, the cell has many ABPs to regulate the kinetics, timing, and precise threshold of how crosslinking will occur. Thus, cytoskeletal polymers may leverage the availability of many ABPs to tune crosslinking and bundling kinetics to set the dynamics of force exertion. The idea that containment of actin in a droplet can lead to feedback between bundling and condensate deformation can have implications in the organization of actin around focal adhesions. Focal adhesion proteins such as talin and vinculin have been shown to form 2D condensate structures on the cellular membrane^10,11,48^. Thus, force generation at the membrane may result from feedback between droplet properties, actin bundling, and membrane properties^49–51^.

While our findings have broad implications for the interactions between the cytoskeleton and protein droplets, we acknowledge that future models require the incorporation of hydrodynamic interactions between the actin filaments and the fluid droplet^52–55^. These hydrodynamic interactions may set the timescales for changes in the curvature of filaments and alter the bending energy of the actin bundles. Other studies involving large numbers of cytoskeletal filaments interacting with motor proteins have established the existence of swirling instabilities in cytoskeletal polymers^56,57^. We anticipate that incorporation of additional physics in our framework has the potential to lead to better understanding of actin responses in liquid-like droplets.

## Author Contributions

D.M., A.C., J.C.S., and P.R. designed the study; D.M., A.C., and C.T.L designed the code and conducted simulations; C.W. and D.J. conducted the experiments. D.M., D.J., C.W., and A.C. analyzed the data. All authors wrote and edited the manuscript.

## Acknowledgments

This research was supported by the National Science Foundation through a MODULUS Grant MCB 2327243 to P.R and J.C.S., and by the National Institutes of Health through R35GM139531 to J.C.S. C.W. was supported through the Center for Dynamics and Control of Materials: an NSF MRSEC under Cooperative Agreement No. DMR-2308817. D.J. was supported by the Ronald E. McNair Scholars Program.

## Supplementary Material for

## Supplementary Methods

### Model Development

#### Chemical and mechanical framework employed in Cytosim

Here, we construct a minimal computational model to explore the emergent properties of actin networks in liquid droplets. Our simulations were performed in Cytosim (https://gitlab.com/f-nedelec/cytosim), an agent-based modeling framework that simulates the chemical dynamics and mechanical properties of cytoskeletal filament networks. Filament dynamics and diffusing species are modeled by numerically solving a constrained Langevin framework in a viscous medium at short time intervals. Additionally, actin filaments are composed of a series of inextensible rigid linear segments of length 0.1 µm that are connected by hinge points to allow for bending. Cytosim computes the bending energy of the fiber using the specified flexural rigidity k_bend_ in the input parameters. Tetrameric VASP crosslinkers are modeled as a single spherical solid of radius 30 nm with four actin-binding sites. Mini-Lpd monomers are modeled as spherical solids of radius 30 nm and function according to the dynamic multimerization model with a single actin-binding site and two monomer-monomer binding sites. Actin binding is governed by rate parameters k_bind_ and k_unbind_. Each binding site has a specified binding distance that specifies the radius of the spherical binding volume within which binding partners are considered as part of the binding reaction. Unbinding reactions are governed by a Bell’s law model representation of slip bond unbinding kinetics where the rate constant is given by the unbinding rate. Steric repulsion potentials between diffusing crosslinkers and filaments are employed to avoid spatial overlap of species.

#### Representation of actin elongation dynamics

Within the existing actin elongation framework, actin filaments elongate deterministically at a constant rate k_grow_. Each filament is represented by a series of segments, each with a maximum length *L*_*seg*_ = 0. 1 µ*m*. Filament elongation rates are scaled to the size of the droplet space such that the filaments grow, if unperturbed, to the length of the circumference of the original droplet, L_fil_ = 2πR µm, by the end of the simulation at 600 s. As such, for a droplet with a radius R = 1 µm where growth is deterministic, each filament will grow at rate k_grow_ = 0.0103 µm/s to reach L_fil_ = 2π µm in length.

#### Dynamics of capping protein

On top of the existing framework for filament elongation dynamics, the effect of capping protein is modeled implicitly by functionalizing the growing end of filaments with the ability to stochastically stop or restart elongation using a similar underlying framework as binding reactions in Cytosim. The capping reaction is governed by the rate parameters k_cap_ and k_uncap_. The implementation of our implicit capping protein model required edits to the Cytosim source code to simulate filament capping and uncapping and have been made available on the GitHub repository alongside this publication.

#### Underlying physics of dynamic ellipsoidal deformation model

The dynamically deformable ellipsoid framework for Cytosim was developed in Dmitrieff et al. 2017^26^ as a computationally expedient method for investigating the role of cortical tension in microtubule assembly within red blood cells. Whereas our previous models used a spherical geometry with a rigid boundary, we now incorporate a deformable ellipsoid geometry that dynamically adjusts the space and axes with each time step to model the condensate surface as a simple continuously deformable surface. Deformation forces are calculated by the force balance between pseudoforces associated with the interfacial surface tension, σ_surface_, the point forces exerted by the filaments on the boundary, and the pseudoforces associated with the pressure^26^. Pressure is represented as a Largrangian multiplier to the constant volume constraint used in this model. Time evolution casts the force balance along the three axes of the ellipsoid and calculates the speed at which deformation occurs as modulated by the effective viscosity, µ_effective_, which acts as a damping parameter to slow the deformation of the ellipse axes over time without affecting the final droplet shape^26^. A full formulation of this model can be found in Dmitrieff et al. 2017^26^.

#### Position evolution

Using the Langevin equation, a stochastic differential equation for describing Brownian motion, Cytosim is able to calculate the time evolution of 3D discretized points for the position of actin filaments and crosslinkers throughout the entire simulation space and duration of the simulation. For a system of N particles, there are 3N coordinates associated with the position of each particle i in 3D space as given by x_i_ = {x_i1_, x_i2_, x_i3_}. Each position x_i_ is then evolved along each dimension j as governed by the following representation of the Langevin equation:

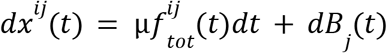

Here, µ is the viscosity of the solvent (not to be confused with µ_effective_). The first term on the right hand side, f^ij^_tot_ (t), is the total force acting on each particle as a function of time. The second term on the right hand side, B_j_(t), is the noise term for diffusion, which is a randomly sampled variable drawn from a normal distribution centered around a mean of 0 and with a standard deviation of 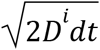. The diffusion constant D is given by the Einstein relation D = µk_B_T where µ is the solvent viscosity, k_B_ is the Boltzmann constant and T is temperature.

### Dynamic multimerization model

We explore the role of multivalent interactions using the dynamic multimerization model where independently diffusing monomers are simulated as solids that each have a single actin-binding site and two multimerization sites. These two multimerization sites allow each monomer to bind with up to two other monomers to dynamically form multimers such as dimers, trimers, tetramers, etc. This model introduces new input parameters that describe the kinetics of multimer formation, k_form_, and multimer splitting, k_split_, and differs from the previous simulations where static crosslinkers are prescribed as a single solid with multiple actin-binding sites. The monomer-monomer binding and unbinding reactions are modeled similarly to actin-crosslinker binding interactions described previously. The implementation of our dynamic multimerization model required edits to the Cytosim source code to simulate multimerization reactions and have previously been made available on a GitHub repository alongside a previous publication^8^.

### Simulations without specific crosslinkers

We explore the role of condensate deformation in directing filament organization in the absence of specific crosslinkers by constructing a model of actin polymerization within a liquid condensate where crosslinkers are omitted and only actin filaments are present within the droplet. This model serves as an analog to droplets composed of RGG, a protein which has only non-specific interactions with actin filaments.

### Theoretical scaling considerations for the droplet diameter and filament thickness

Here, we present the salient arguments laid out in Limozin et al.^5^ and rederived in our earlier study^9^. Consider a liquid droplet of radius R with an actin bundle of length L. Actin bundle is present below the interface between droplet and the surrounding medium. We assume that the thermal fluctuations of the bundle dominate the droplet shape. The coordinate of the bundle centerline subject to fluctuations can be written as,

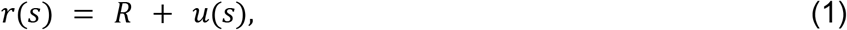

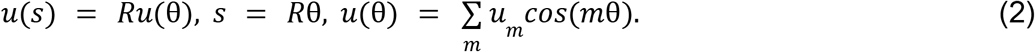

Here, *s* represents arc length, *r*(*s*) represents the radial coordinate of bundle centerline as a function of arc length. The radial coordinate is represented fluctuations *u*(*s*) around the droplet radius.

The bending energy of the bundle with flexural rigidity *k* _*b*_ with local curvature κ(*s*) is given by,

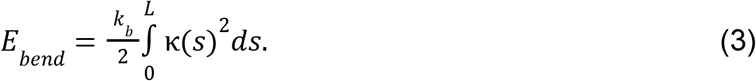

The flexural rigidity is related to persistence length *L*_*p*_ of the bundle by, *k*_*b*_ = *L*_*p*_ *k*_*B*_ *T*, where T is temperature and *k*_*B*_ is the Boltzmann constant. The local curvature can be written in polar coordinate as,

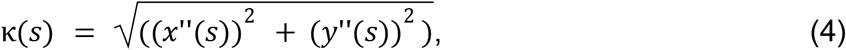

where,

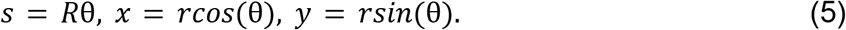

Substituting, we get,

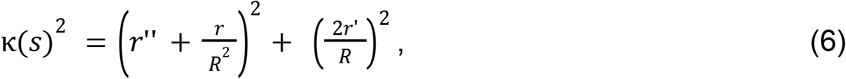

where,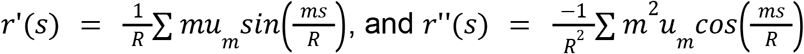.

Now, we assume that the different modes (m with amplitude *u*_*m*_) are uncorrelated, and ignore cross correlations to write⟨κ(*s*)^2^ ⟩,

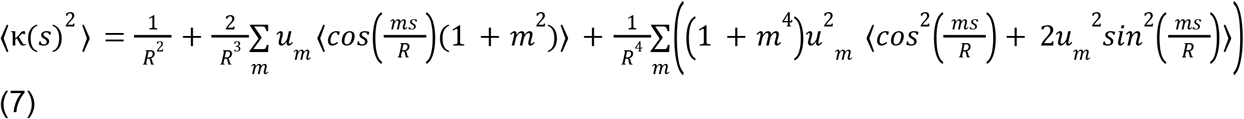

Please note that the primary contribution to ⟨κ(*s*) ^2^ ⟩ comes from the global curvature (droplet curvature) 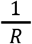 while the fluctuations contribute to higher order terms.

Substituting Eq. (7) in (3), we get,

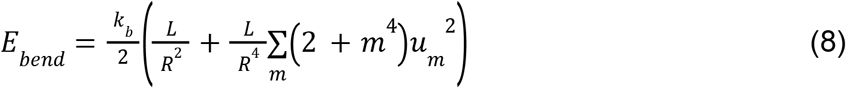

The primary contribution to the bending energy comes from droplet radius. The fluctuations above that baseline energy, Δ*E*_*bend*_ is given by,

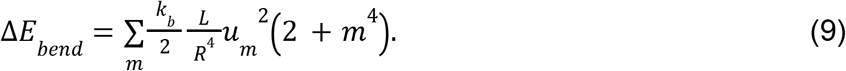

Applying equipartition theorem, each mode contributes *k*_*B*_ *T*/2 energy, where *k*_*B*_ is the Boltzmann constant and temperature *T* as given by,

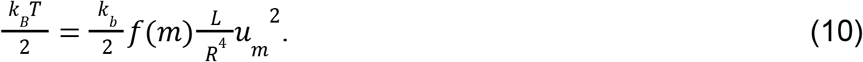

Hare, we represent the mode-dependent terms by the function *f*(*m*). Assuming that the ring thickness is proportional to the Fourier amplitude *u*_*m*_, for a bundle of length *L* = 2π*R*, we get,

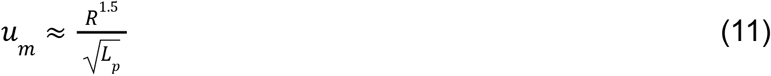

Through this, we show that the bundle thickness has a power law relationship with droplet radius.

## Experimental Methods

### Reagents

Tris base, NaCl, Hellmanex III, Tris(2-carboxyethyl)phosphine (TCEP), poly-L-lysine, Atto 594 maleimide, Atto 488 maleimide, were purchased from Sigma-Aldrich. mPEG SVA was purchased from Laysan Bio. Phalloidin-iFluor594 was purchased from Abcam. Rabbit muscle actin was purchased from Cytoskeleton. Capping Protein (CapZ) was purchased from Hypermol.

### Plasmids

A pET vector encoding the ‘cysteine light’ variant of human VASP (pET-6xHis-TEV-KCK-VASP(CCC-SSA)) and monomer mini-Lpd (his-EGFP-Lpd(aa850-1250) were gifts from Scott Hansen.

### Protein Purification

The mini-Lpd (his-Z-EGFP-LZ-Lpd(aa850-1250)) was transformed into BL21 (NEB, cat. No. C2527H) and grown at 30 °C to an OD of 0.8. The bacteria were then cooled to 12°C and induced for 24 hours with 1mM IPTG. The rest of the protocol was performed at 4 °C. Cells were pelleted from a 2L culture by centrifugation at 4,785g (5,000 rpm in Beckman JLA-8.100) for 20 min. Pellets were resuspended in 100mL of lysis buffer (50mM sodium phosphate pH 8.0, 300mM NaCl, 10mM imidazole, 0.5mM TCEP, 0.2% Triton X100, 10% glycerol, 1mM PMSF, and EDTA free protease inhibitor tablets (1 tablet per 50mL) (Roche cat# 05056489001)) followed by sonication on ice for 4×2000J with amplitude at 10 (Sonicator Qsonica LLC, Q700). The lysate was clarified by centrifugation at 48,384g (20,000 rpm in Beckman JA25.50) for 30 min at 4 °C before being applied to a 10mL bed volume Nickel nitrilotriacetic acid (Ni-NTA) agarose (Qiagen, cat. no. 30230) column, and washed with 10 column volumes (CVs) of lysis buffer to which imidazole had been added to a final concentration of 20mM. The column was then washed with 5 column volumes of lysis buffer containing 20 mM Imidazole but lacking Triton-X100 and protease inhibitor tablets. The protein was eluted with elution buffer (50 mM Tris-HCl, pH 7.5, 300mM NaCl, 10% glycerol, 400mM imidazole, 1mM TECP, 1mM PMSF, and EDTA-free protease inhibitor tablets (1 tablet per 50mL). The protein was concentrated using Amicon Ultra-15, 30K MWCO (Millipore: Cat#UFC903024) to 5 mL, and clarified by ultracentrifugation for 5min at 68,000 x g (40,000 rpm with Beckman optimal MAX-E Ultracentrifuge and TLA100.3 rotor). The protein was further purified by size exclusion chromatography with Superose 6, and ion exchange chromatography with SP Sepharose Fast Flow (GE Healthcare, Cat#17-0729-01), and stored as liquid nitrogen pellets at -80°C.

The pET-His-KCK-VASP(CCC-SSA) plasmid was transformed into Escherichia coli BL21(DE3) competent cells (NEB, cat. no. C2527). Cells were grown at 30 °C to an optical density (OD) of 0.8. Protein expression was performed as described previously with some alteration^9^. Expression of VASP was induced with 0.5 mM isopropylthiogalactoside (IPTG), and cells were shaken at 200 rpm at 12 °C for 24 h. The rest of the protocol was carried out at 4 °C. Cells were pelleted from 2 L cultures by centrifugation at 4,785g (5,000 rpm in Beckman JLA-8.100) for 20 min. Cells were resuspended in 100 mL lysis buffer (50 mM sodium phosphate pH 8.0, 300 mM NaCl, 5% glycerol, 0.5 mM TCEP, 10 mM imidazole, 1 mM phenylmethyl sulphonyl fluoride (PMSF)) plus EDTA-free protease inhibitor tablets (1 tablet per 50 mL, Roche, cat. no. 05056489001), 0.5% Triton-X100, followed by homogenization with a dounce homogenizer and sonication (4 × 2,000 J). The lysate was clarified by ultracentrifugation at 125,171g (40,000 rpm in Beckman Ti45) for 30 min. The clarified lysate was then applied to a 10 mL bed volume Nickel nitrilotriacetic acid (Ni-NTA) agarose (Qiagen, cat. no. 30230) column, washed with 10 column volumes of lysis buffer plus EDTA-free protease inhibitor tablets (1 tablet per 50 mL), 20 mM imidazole, 0.2% Triton X-100, followed by washing with 5×CV of lysis buffer plus 20 mM imidazole. The protein was eluted with elution buffer (50 mM Tris, pH 8.0, 300 mM NaCl, 5% glycerol, 250 mM imidazole, 0.5 mM TECP, EDTA-free protease inhibitor tablets (1 tablet per 50 mL)). The protein was further purified by size exclusion chromatography with Superose 6 resin. The resulting purified KCK-VASP was eluted in storage buffer (25 mM HEPES pH 7.5, 200 mM NaCl, 5% glycerol, 1 mM EDTA, 5 mM DTT). Single-use aliquots were flash-frozen using liquid nitrogen and stored at −80 °C until the day of an experiment.

Expression and purification of his-LAF-1 RGG, containing amino acids 1-168 of LAF-1 and a C-terminal 6xHis tag, was performed as described previously, including starting plasmids^58^. In brief, RGG was overexpressed in E. coli BL21(DE3) cells. One-liter cultures were induced with 0.5 mM IPTG overnight at 18C and 220 rpm, and pellets of cells expressing his-LAF-1 RGG were harvested from the cultures via centrifugation when the OD 600 reached 0.8. The pellets were then lysed in a buffer containing 20 mM Tris, 500 mM NaCl, 20 mM imidazole, and one EDTA-free protease inhibitor tablet (SigmaAldrich) for 5 min on ice and then sonicated. The cell lysates were centrifuged at 134,000 g for 40 min, after which his-LAF-1 RGG resided in the supernatant. The supernatant was then mixed with Ni-NTA resin (G-Biosciences) for 1 h, settled in a glass column, and washed with a buffer containing 20 mM Tris, 500 mM NaCl, 20 mM imidazole (pH 7.5). The bound proteins were eluted from the Ni-NTA resin with a buffer containing 20 mM Tris, 500 mM NaCl, 500 mM imidazole (pH 7.5). The purified proteins were then buffer exchanged into 20 mM Tris, 500 mM NaCl (pH 7.5) using Amicon Ultra centrifugal filters. Small aliquots of the protein were frozen in liquid nitrogen at a protein concentration of approximately 120 mM and stored at 80C. To promote solubility of the LAF-1 RGG protein, the entire purification process was performed at room temperature, except for the cell lysis process, which was done on ice.

### Protein Labeling

The VASP used in this study is a previously published ‘cysteine light’ mutant that replaced the three endogenous cysteines with two serines and an alanine. A single cysteine was then introduced at the N-terminus of the protein to allow selective labeling with maleimide dyes. This mutant was found to function in an indistinguishable manner from the wild-type proteins^59^. Thus, VASP was labeled at the N-terminal cysteine using maleimide-conjugated dyes. VASP was buffer exchanged into 20 mM Tris (pH 7.4) 150 mM NaCl buffer to remove DTT from the storage buffer and then incubated with dye for two hours at room temperature. Free dye was then removed by applying the labeling reaction to a spin column packed with Sephadex G-50 Fine DNA Grade (GE Healthcare GE17-0573-01) hydrated with TNT buffer (20 mM Tris pH 7.4, 150 mM NaCl, and 5 mM TCEP). The labeled protein was then centrifuged at 100,000 x G for 10 min at 4 degrees Celsius to remove aggregates before being flash-frozen in single-use aliquots.

Monomeric actin was labeled using maleimide-conjugated dyes. Dyes were incubated with G-actin for 2 hours at room temperature before being separated from the labeled protein by applying the labeling reaction to a spin column packed with Sephadex G-50 Fine DNA Grade (GE Healthcare GE17-0573-01) hydrated with A buffer (5 mM Tris-HCL (pH 8), 0.2 mM ATP and 0.5 mM DTT pH 8). The labeled protein was then centrifuged at 100,000 x G for 10 min at 4 degrees Celsius to remove aggregates before being flash-frozen in single-use aliquots.

RGG was labeled using amine-reactive NHS Ester dyes. RGG was buffer exchanged into 25mM HEPES (pH 7.4) 500mM NaCl prior to labeling with dyes. After labeling RGG was separated from unconjugated dye and buffer exchanged back to 20 mM Tris (pH 7.4) 500mM NaCl buffer using 3K Amicon columns (MilliporeSigma, Burlington, MA). Small aliquots of the protein were frozen in liquid nitrogen and stored at −80°C.

### Protein droplet formation and actin polymerization

VASP, monomer mini-Lpd, or RGG droplets were formed by mixing the respective proteins with 3% (w/v) PEG-8000 in 20 mM Tris pH 7.4, 5 mM TCEP, and either 150 mM NaCL (for VASP) or 50 mM NaCl (for monomer mini-Lpd and RGG). PEG was added last to induce droplet formation. All protein concentrations refer to the monomeric concentrations. For actin polymerization assays within protein droplets, protein droplets were allowed to form for ten minutes following PEG addition following which monomeric actin was mixed into the droplet solution. Atto-594-labelled G-actin was allowed to polymerize in the droplets for 15 min, then the droplets were imaged. For phalloidin-actin assays unlabelled actin monomers were added to protein droplets and allowed to polymerize for 15 min. Phalloidin-iFluor594 (Abcam) was then added and allowed to stain filamentous actin for 15 min prior to imaging.

### Microscopy

Samples were prepared for microscopy in 3.5mm or 5mm diameter wells formed using biopsy punches to create holes in 1.6 mm thick silicone gaskets (Grace Biolabs) on Hellmanex III cleaned, no. 1.5 glass coverslips (VWR). Coverslips were passivated using poly-L-lysine conjugated PEG chains (PLL-PEG). To prevent evaporation during imaging, an additional small coverslip was placed on top of the gasket to seal the well. Fluorescence microscopy was done using an Olympus SpinSR10 spinning disk confocal microscope with a Hamamatsu Orca Flash 4.0V3 Scientific CMOS camera. FRAP was done using the Olympus FRAP unit 405 nm laser.

PLL-PEG was prepared as described previously with minor alterations^60^. Briefly, amine-reactive mPEG succinimidyl valerate was conjugated to poly-L-lysine at a molar ratio of 1:5 PEG to PLL. The conjugation reaction takes place in 50mM sodium tetraborate solution pH 8.5 and is allowed to react overnight at room temperature while continuously stirring. The final product is then buffer exchanged to PBS pH 7.4 using 7000 MWCO Zeba spin desalting columns (Thermo Fisher) and stored at 4 °C.

### Image Analysis

Image J was used to quantify the distribution of condensate characteristics. Specifically, condensates were selected using thresholding in the brightest channel and shape descriptors (i.e. diameter, aspect ratio, etc.), and protein fluorescent intensities were measured using the built-in analyze particles function. For aspect ratio analysis condensates that had come into contact with other condensates were removed from the analysis to avoid any skewing of data from misrepresentation of single condensate deformation.

For actin ring thickness and power law fitting analysis quantification of the ring thickness within the protein condensates was performed by measuring the phalloidin-actin intensity across a radial line bisecting the ring, taking the full-width at half-maximum intensity of the actin ring using a gaussian fit of the intensity profile across one side of the actin ring, and then subtracting the diffraction limit to get an approximate ring thickness. To compare droplet diameters among aspherical droplets with AR > 1, we calculated an effective spherical diameter for the droplets based on their major and minor axis lengths: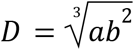, where a refers to the major axis and b refers to the minor axis lengths.

## Data Analysis

### Filament length probability density function calculation

The lengths of each filament is calculated by adding up the length of each segment at each time step. A filament length distribution is then created by sorting the lengths into 130 bins of equal width 0.05 µm. Since the filament length distribution changes with each time step, we then smooth the distribution over time by calculating a moving average of filament lengths over a period of ±5 time steps (±5 s) at each time step. Then we feed this into a kernel density estimation to form a Gaussian approximation of the probability density function of filament lengths at each timestep.

### Actin-covered surface fraction calculation

As the actin filaments grow to form shells, discs, and rings, they will move to occupy space near the surface of the droplet. Since the condensate boundary is modeled as a dynamically deformable ellipsoid, the droplet shape may be different at each time point. As such, at each time point, we discretize the surface of the droplet to an icosphere that is then deformed according to the current state of the ellipsoid. Actin filaments are also discretized to the monomer level and used to calculate the surface density at each time point via the fraction of occupied triangles using a threshold distance of 0.1 µm from the surface of the deformed icosphere. More details about the method can be found in our previous study^18^.

### Filament alignment angle calculation

Throughout the duration of our simulations, actin elongation within the confinement of a droplet drives the rearrangement of individual filaments which leads to the emergence of actin network structures such as shells, discs, and rings. These structures differ in how filaments are organized with respect to one another. As such, we define an alignment angle as a metric that describes the extent to which the actin network structure has collapsed from a 3D spherical shell to a tight 2D ring, inclusive of intermediary shapes. At each time point, we first discretize the actin filaments to the monomer level and use the resulting filament positions to construct the gyration tensor for the filament network. We next extract the eigenvalues of the gyration tensor to compute the ellipsoidal spans, thus providing us with the axis lengths for an ellipsoidal shape approximation that best explains the distribution of filaments in the network. The alignment angle is thus defined as the angle between the major axis (span a) and minor axis (span c) of the ellipsoidal approximation. As such, the filament alignment angle varies from 0 degrees (perfectly flat) to 45 degrees (perfectly spherical).

## Supplementary Tables

**Table S1:** Table of universal parameters used in the Cytosim model.

| Parameter | Value | Notes/Reference |
| --- | --- | --- |
| Total time ($T_{sim}$) | 600 s | |
| Time step | 0.002 s |  |
| Condensate viscosity | 0.5 pN s/$\mu\text{m}^2$ | 500x water;<br>Chosen based on protein condensate viscosities<sup>61</sup>. |
| **Boundary** |  |  |
| Initial Shape | Spherical, constrained to a deformable ellipsoid unless stated otherwise. | Rigid Ellipsoidal in **Fig. 8A**<br>Rigid Spherical in **Fig. S1** |
| Effective Viscosity: $\mu_{effective}$ | 100 pN s/$\mu\text{m}$ unless stated otherwise. | Varied in **Fig. 7** and **Fig. S3** to show that this parameter does not affect droplet deformation timescales in our study |
| Radius | 1 $\mu\text{m}$ unless stated otherwise. | Varied in power law simulations. |
| Boundary repulsion stiffness | 200 pN/$\mu\text{m}$ for actin filaments;<br>100 pN/$\mu\text{m}$ for crosslinking molecules | This specifies the spring stiffness that acts on the discretized points of each actin filament and crosslinking molecule if the point lies outside the specified boundary. The force on each point is dependent on the distance that it lies beyond the confines of the boundary. |
| **Actin filaments** |  |  |
| Segmentation length $L_{seg}$ | 0.1 $\mu\text{m}$ (100 nm) | |
| Maximum length | $2\pi R$ $\mu\text{m}$ | |
| Polymerization rate $k_{grow}$ | 0.0103 $\mu\text{m/s}$ for diameter 2 $\mu\text{m}$<br>General formula = $(2\pi R - L_{fil}^0)/T_{sim}$ | Only plus (+) end extension is allowed. This rate is calculated by assuming a final filament length of $2\pi$ $\mu\text{m}$ at 600 s. |
| Brownian ratchet force for polymerization | 10 pN | 43 |
| Actin flexural rigidity $k_{bend}$ | 0.075 pN$\square\mu\text{m}^2$ | 25 |
| Actin steric repulsion $k_{\text{steric}}$ | Radius 3.5 nm<br>Stiffness 1.0 pN/$\mu\text{m}$ | Chosen to ensure the observation of ring structures within the kinetic parameters used in this study as determined from a previous study<sup>18</sup>. |
| **Tetravalent crosslinkers (VASP)** |  |  |
| Radius | 30 nm |  |
| Diffusion rate | 10 $\mu\text{m}^2/\text{s}$ | |
| Concentration of tetramers | 0.40 $\mu\text{M}$ [1000 tetramers when Radius = 1 $\mu\text{m}$] | The number of simulated tetramers is scaled to maintain the equivalent concentration with varied droplet size. |
| Actin-binding rate (Ring Kinetics) | 10.0 (1/s) | Determined from VASP simulations. |
| Actin-binding rate (Shell Kinetics) | 0.1 (1/s) | Determined from VASP simulations. |
| Actin-binding distance | 30 nm |  |
| Actin-binding valency | 4 | Each spherical molecule approximates a VASP tetramer. |
| Zero-force actin-unbinding rate (Ring and Shell Kinetics) | 1.0 (1/s) | Determined from VASP simulations. |
| Actin-unbinding force | 10 pN | Typical values for passive crosslinkers<sup>62</sup>. |
| VASP steric repulsion | Radius 30 nm<br>Stiffness 10 pN/$\mu\text{m}$ | Chosen to ensure the observation of ring structures within the kinetic parameters used in this study as determined from a previous study on tetramers<sup>18</sup>. |
| **Monovalent actin binders that interact with each other multivalently (mini-Lpd monomers)** |  |  |
| Radius | 30 nm |  |
| Diffusion rate | 10 $\mu\text{m}^2/\text{s}$ | |
| Concentration of monomers | 0.79 $\mu\text{M}$ [2000 monomers when radius = 1 $\mu\text{m}$] | The number of simulated monomers is scaled to maintain the equivalent concentration with varied droplet size. |
| Actin-binding rate | 10.0 (1/s) | Determined from VASP simulations. |
| Actin-binding distance | 30 nm |  |
| Actin-binding valency | 1 | Each spherical molecule approximates a mini-Lpd monomer. |
| Zero-force actin-unbinding rate | 1.0 (1/s) | Determined from VASP and mini-Lpd simulations. |
| Actin-unbinding force | 10 pN | Typical values for passive crosslinkers<sup>62</sup>. |
| mini-Lpd monomer-actin steric repulsion | Radius $R_{\text{solid}} = 30$ nm<br>Stiffness 10 pN/$\mu\text{m}$ | Chosen to ensure the observation of ring structures within the kinetic parameters used in this study as determined from a previous study on tetramers and mini-Lpd monomers<sup>8,18</sup>. |
| mini-Lpd monomer-monomer binding distance | 90 nm ($3R_{\text{solid}}$) | This is the distance between the binding site on solid A and the center of solid B. So, if two solids are in contact, depending on the position of the binding site this distance can scale between $R_{\text{solid}}$ and $3R_{\text{solid}}$, where $R_{\text{solid}}$ is the radius of solid. |
| Monomer-monomer binding valency | 2 | Chosen to allow for the formation of higher multimeric states. |
| mini-Lpd monomer-monomer splitting force | 10 pN | Used in this study |
| mini-Lpd monomer-monomer steric repulsion | Radius 30 nm<br>Stiffness 5.0 pN/$\mu\text{m}$ | Chosen empirically to ensure adequate dimerization reactions occur to support the observation of ring structures as determined from a previous study on mini-Lpd monomers<sup>8</sup>. |

**Table S2:**
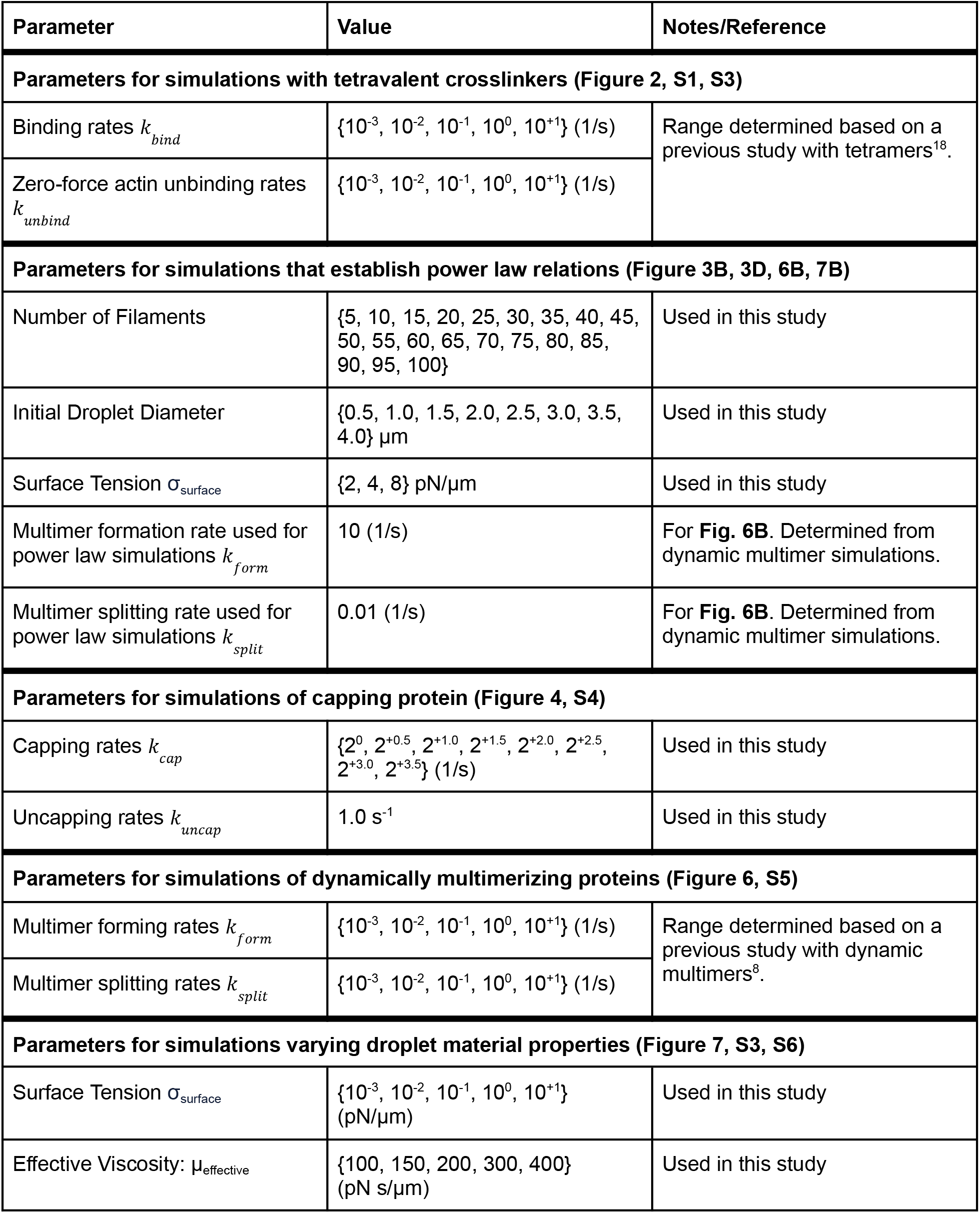
Table of varied parameters.

## Supplementary Figures

**Figure S1:**
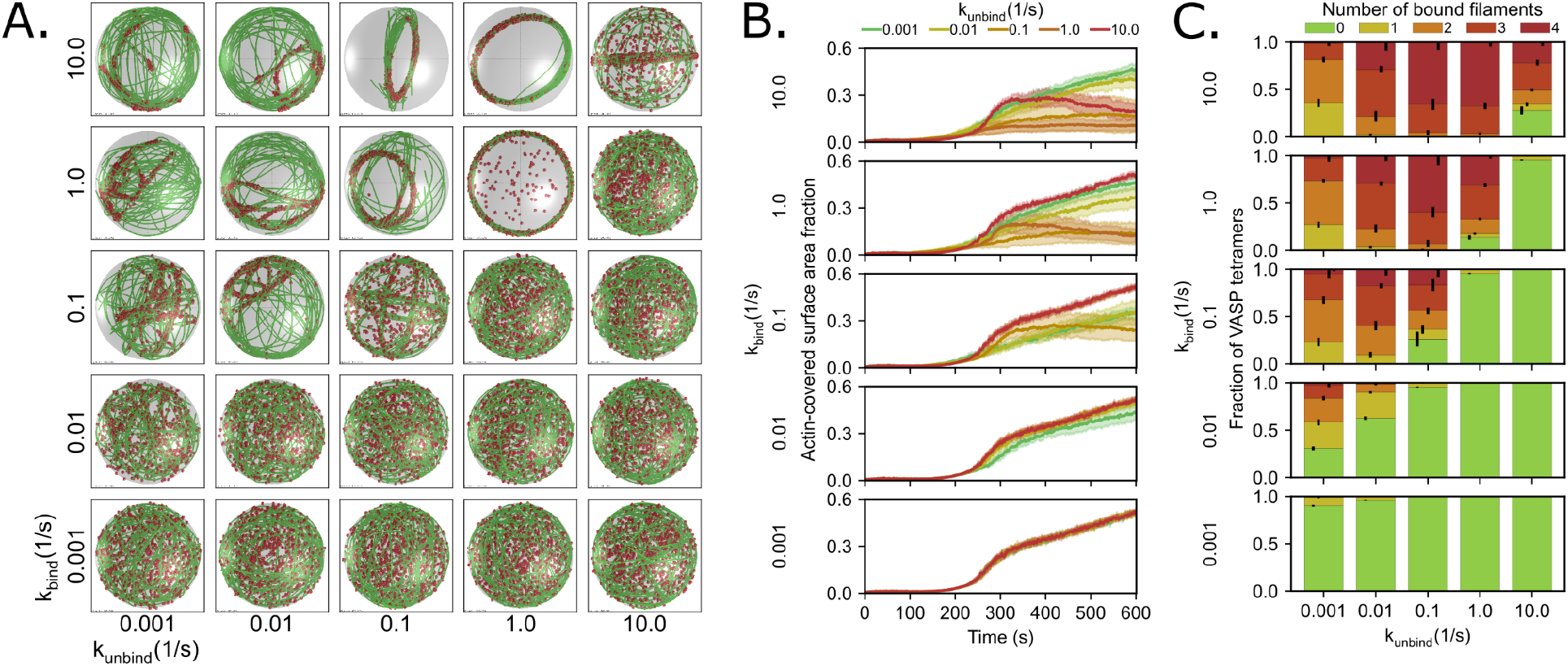
Varied crosslinker kinetics in simulations of rigid-boundary spherical condensates. **A)** Representative final snapshots (t = 600 s) from simulations at various binding and unbinding rates within rigid-boundary spherical condensates (R = 1 μm) containing 30 actin filaments (green) and 1000 tetravalent crosslinkers (red spheres). The binding rates of the tetravalent crosslinkers are varied along each column, and unbinding rates are varied along each row. The polymerization rate at the plus (+) end is constant at 0.0103 μm/s, and neither end undergoes depolymerization. **B)** Actin-covered surface area fraction for varied tetravalent crosslinker binding and unbinding kinetics. **C)** Fraction of tetravalent crosslinkers bound to 0, 1, 2, 3, or 4 actin filaments for each condition. The error bars represent the standard deviation. Data was obtained from the last 30 snapshots (5%) of each replicate. For **B** and **C**, 10 replicates are considered per condition.

**Figure S2:**
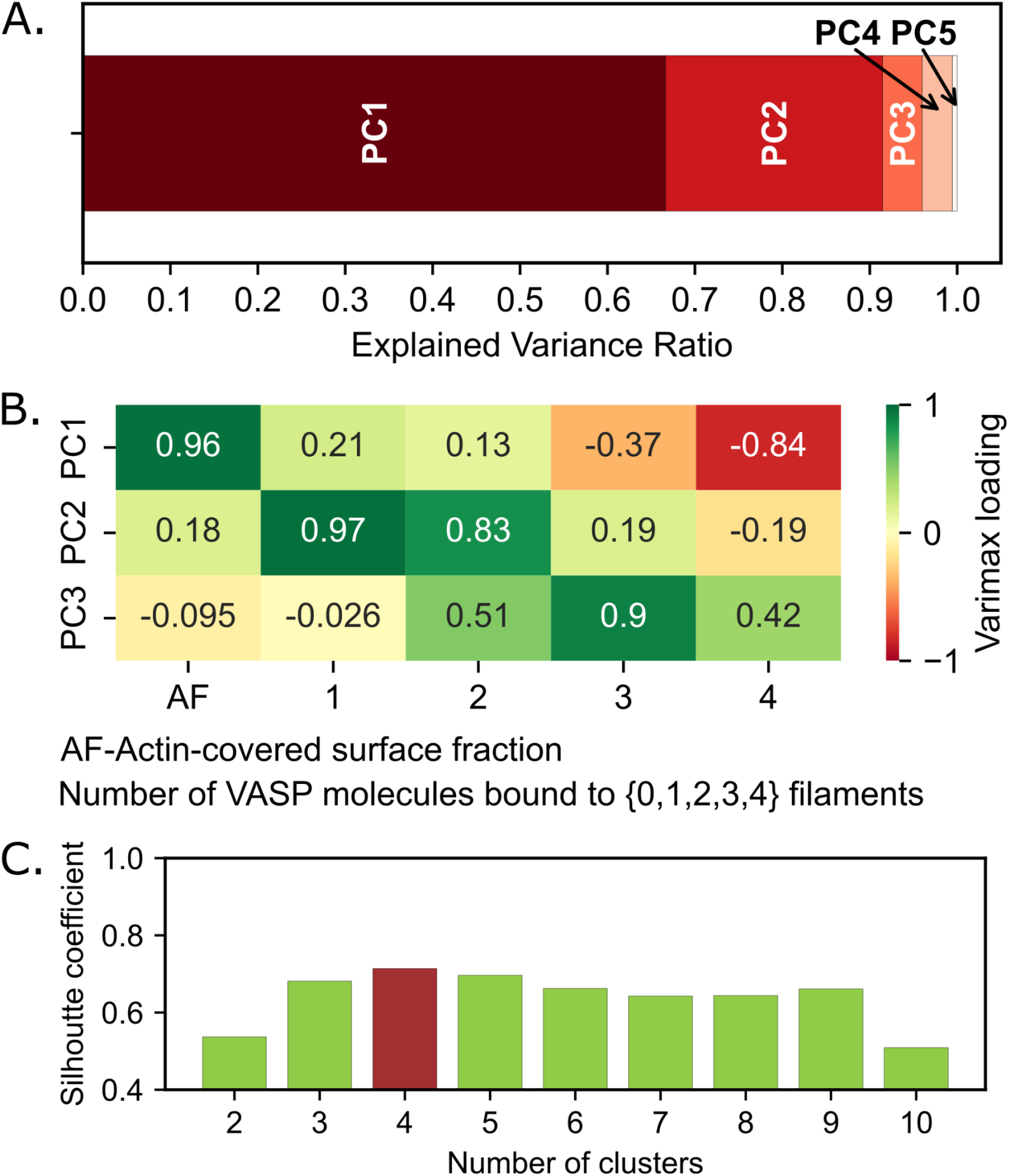
K-means clustering and PCA optimization. **A)** Principal component analysis was employed to find orthogonal axes corresponding to the data represented by the actin-covered surface fraction and the fraction of VASP tetramers bound to 1, 2, 3, and 4 actin filaments. **B)** Varimax loadings were calculated for each of the PCs with the five factors listed above. Loadings reveal that PC1 primarily reflects information positively correlated with the actin-covered surface fraction and negatively correlated with the fraction of VASP tetramers bound to four filaments, PC2 reflects a positive correlation with the fraction of VASP bound to one and two filaments, while PC3 reflects a positive correlation with the fraction of VASP bound to 3 filaments. **C)** Silhouette coefficient shows that the dataset is made up of a maximum of 4 clusters. Data used: Last 5 snapshots from each of the 10 replicates per (*k bind, k*_*unbind*_) pair value.

**Figure S3:**
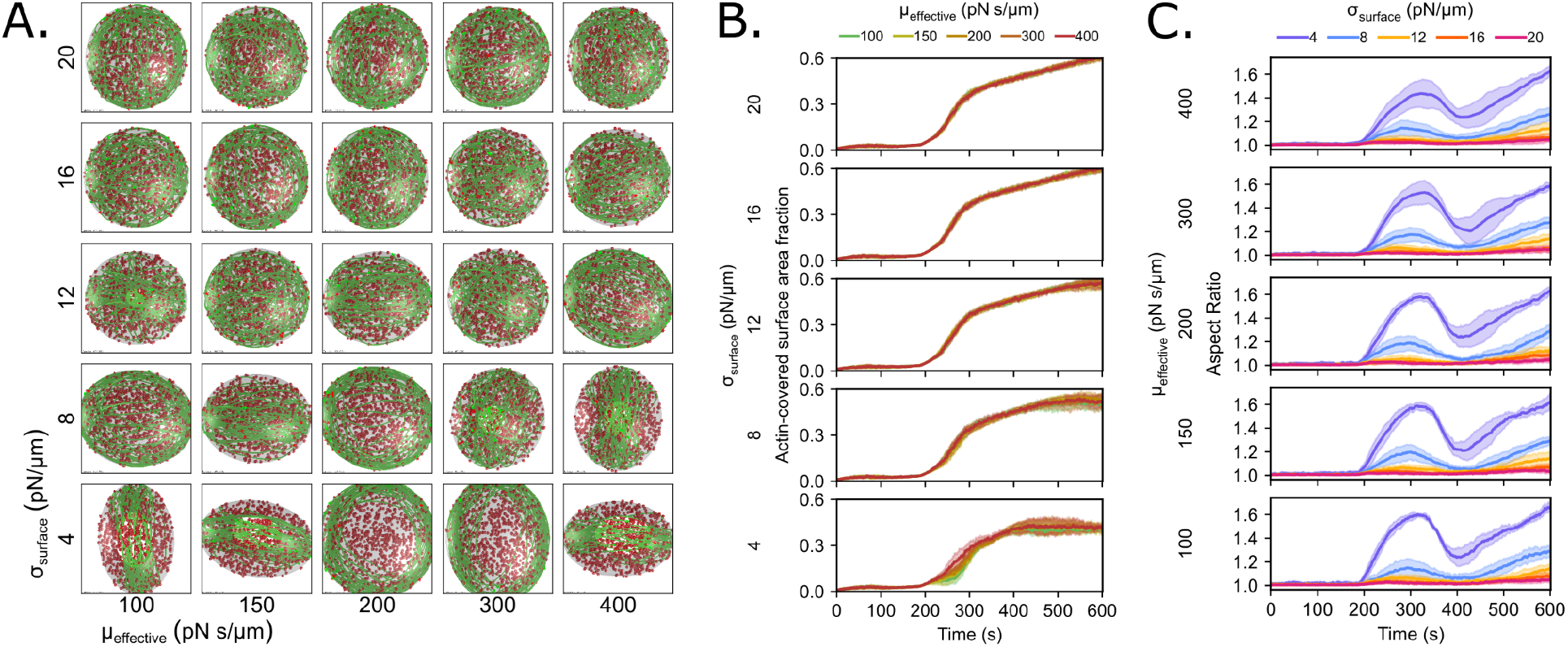
Interfacial properties of the condensate surface govern deformation and determine whether the actin network will form shells or discs. **A)** Representative final snapshots (t = 600 s) from simulations varying the properties of the deformable condensate boundary (initially spherical with R = 1 μm) containing 30 actin filaments (green) and 1000 tetravalent crosslinkers (red spheres) with shell/disc-forming kinetics. The effective viscosity which attenuates the rate of deformation is varied along each column, and the surface tension which describes the innate resistance of the condensate boundary to deformation is varied along each row. The polymerization rate at the plus (+) end is constant at 0.0103 μm/s, and neither end undergoes depolymerization. **B)** Time series showing the mean (solid line) and standard deviation (shaded area) of the actin-covered surface area fraction for varied condensate interfacial properties. **C)** Time series showing the mean (solid line) and standard deviation (shaded area) of condensate aspect ratio for each condition. Surface tension is varied within each plot with the same effective viscosity. The aspect ratio is defined as the ratio between the longest and shortest axis of the ellipsoid (AR = a/c) where a ≥ b ≥ c.

**Figure S4:**
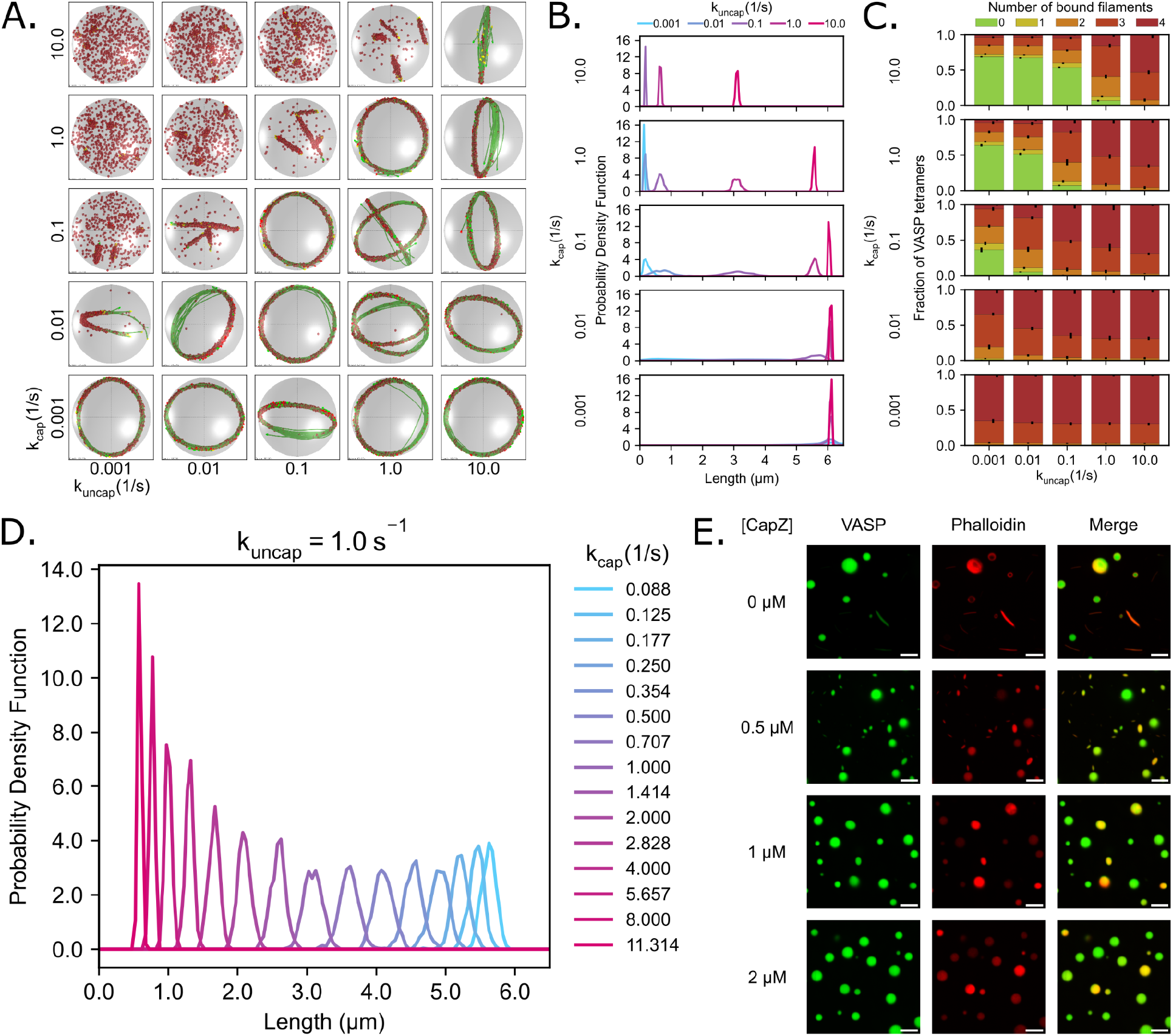
Capping and uncapping kinetics of capping protein tunes filament length in rigid condensates. **A)** Representative final snapshots (t = 600 s) from 10 replicates at various capping and uncapping rates within condensates with a rigid spherical boundary (R = 1 μm) containing 30 actin filaments (green) and 1000 tetravalent crosslinkers (red spheres). The filament growth rate is fixed at 0.0103 μm/s, and the capping rates are varied along each column, and the uncapping rates are varied along each row. The binding rates of the tetravalent crosslinkers are chosen to promote ring formation when L_fil_ = 2πR. **B)** Probability density function of the filament length for each simulation condition shown in **A**. As the capping rate increases, the probability of finding longer filaments decreases. The uncapping rate k_uncap_ is changed within each subpanel and the capping rate k_cap_ is changed between the subpanels. **C)** Stacked bar graph shows the mean fraction of tetravalent crosslinkers bound to 0, 1, 2, 3, or 4 actin filaments for each condition. The error bars represent the standard deviation. The data used is from the last 30 snapshots (5%) of each of the 10 replicates. **D)** Probability density function of the filament length for an extended set of simulation conditions where capping rates are varied while the uncapping rate is held constant at 1.0 s^-1^ to sample the transition between short rods and ring structures within deformable condensates. The deformable boundary has a surface tension of 4 pN/µm and an effective viscosity of 100 pN s/µm. As the capping rate increases, the probability of finding longer filaments decreases. Additionally, the probability density function narrows when k_uncap_/k_cap_ is small or large and is broader when k_cap_ ≈ k_uncap_. **E)** Phalloidin-iFluor-594 staining of Atto 488 labeled VASP condensates with 3 μM actin, displaying polymerized actin within the condensates and the disruption of rings with increasing CapZ (not labelled) concentration. Scale bars, 5 μm.

**Figure S5:**
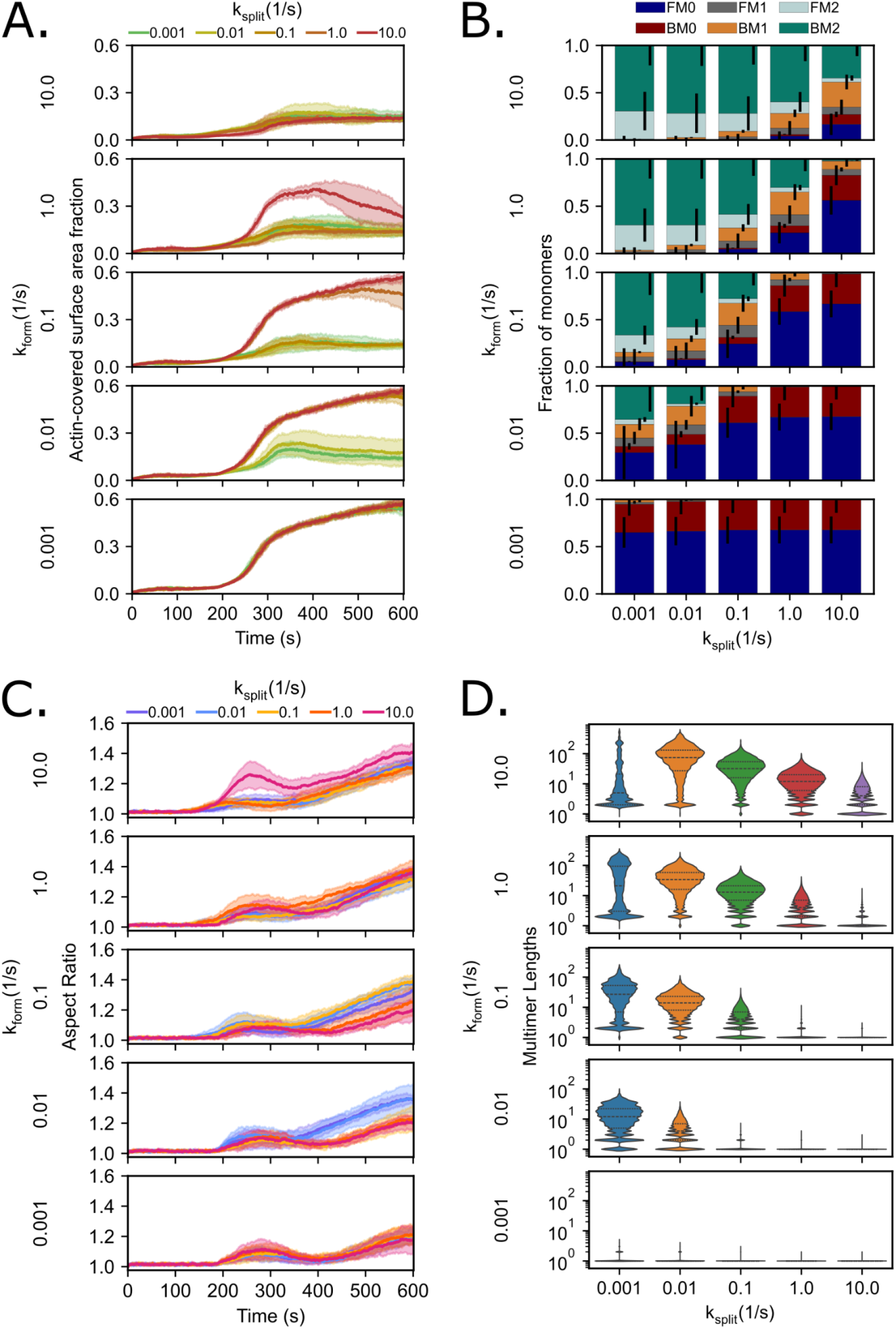
Multivalent condensate environments exhibit disc formation and a power law scaling relationship between the initial diameter of the condensate and the number of filaments. **A)** Time series showing the mean (solid line) and standard deviation (shaded area) of the actin-covered surface area fraction for varied multimer forming and splitting kinetics. **B)** Stacked bar graph shows mean fraction of monomers in various multimerization and actin-bound states for each condition. BM (Bound Monomer) refers to monomeric units that are bound to an actin filament, while FM (Free Monomer) refers to those that are not bound to actin; the corresponding number (0, 1, or 2) indicates the number of other monomers that a single monomeric unit is bound to. The error bars represent the standard deviation. Data was obtained from the last 30 snapshots (5%) of each replicate. **C)** Time series showing the mean (solid line) and standard deviation (shaded area) of condensate aspect ratio for each condition. The aspect ratio is defined as the ratio between the longest and shortest axes of the ellipsoid (AR = a/c), where a ≥ b ≥ c. **D)** Violin plots showing the distribution of multimer lengths for each simulation condition for systems. Lengths are counted as the number of monomers that constitute a single multimer chain. Violin plot densities are normalized such that all plots are fit to the same width. Median (dashed line), and quartiles (dotted lines) are shown. Data was obtained from the last 30 snapshots (5%) of each replicate. For **A, B, C**, and **D**, 10 replicates are considered per condition.

**Figure S6:**
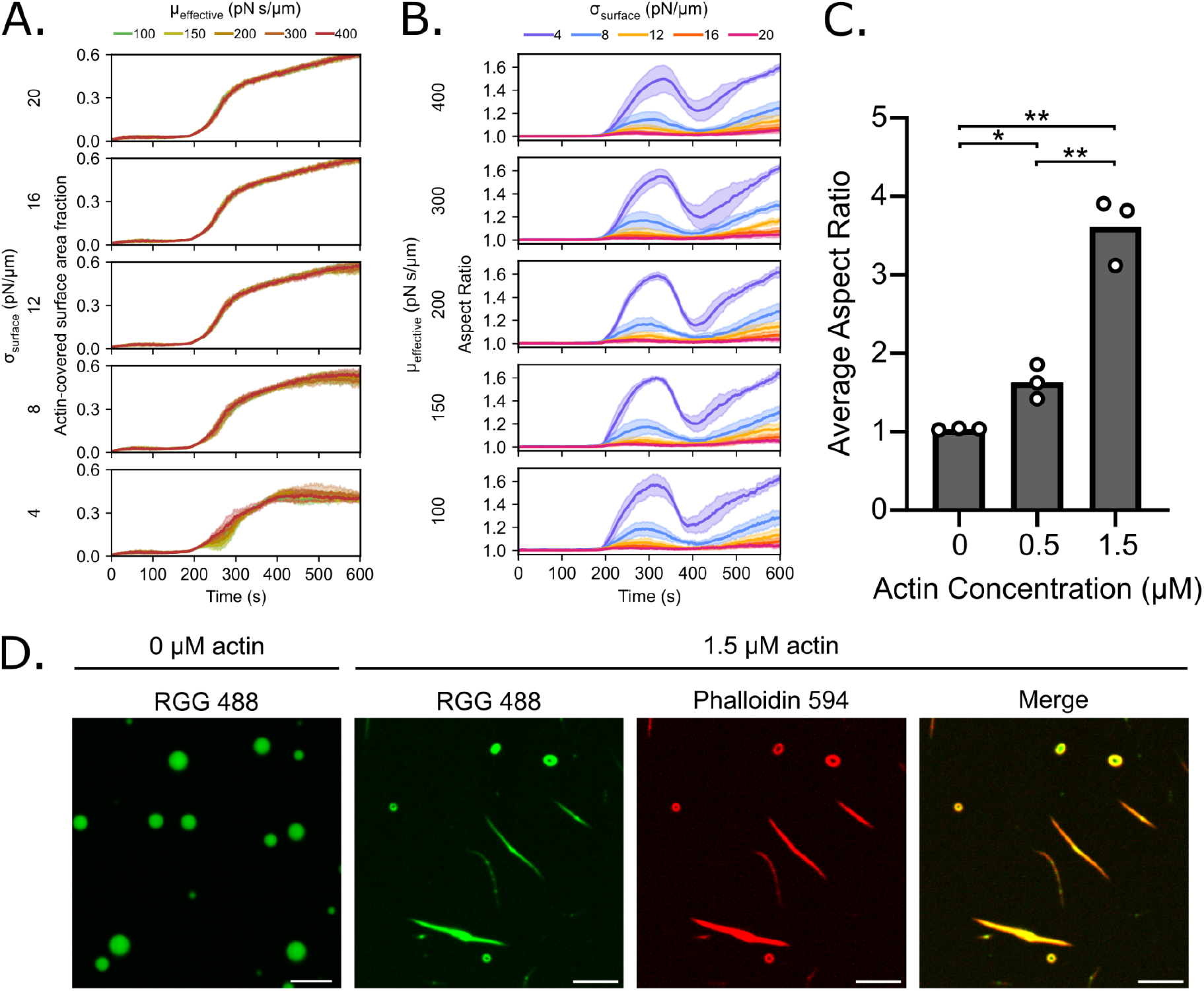
Interfacial properties of the condensate surface govern deformation and determine whether the actin network will form shells or discs. **A)** Each subpanel shows the time series of mean (solid line) and standard deviation (shaded area) of the actin-covered surface area fraction colored by effective viscosity at a given surface tension value mentioned to the left. **B)** Each subpanel shows the time series of mean (solid line) and standard deviation (shaded area) corresponding to condensate aspect ratio colored by surface tension (value shown in legend above). The effective viscosity of the droplet interface is varied between the subpanels and is mentioned to the left. Surface tension is varied within each plot with the same effective viscosity. The aspect ratio is defined as the ratio between the longest and shortest axis of the ellipsoid (AR = a/c) where a ≥ b ≥ c. For **A** and **B**, 10 replicates are considered per condition. **C)** Quantification of the average aspect ratio for conditions shown in **7C**, showing an increase in aspect ratio with increasing actin concentration. Data are the mean across three independent experiments. The overlaid white circles denote the means of each replicate. One asterisk denotes p <.05, two asterisks denote p <.01, and three asterisks denote p <.001 using an unpaired, two-tailed t-test on the means of the replicates, n=3. Buffer conditions for all conditions were 20 mM Tris pH 7.4, 50 mM NaCl, 5 mM TCEP, and 3% (w/v) PEG 8000. D) Phalloidin-iFluor-594 staining of Atto 488 labeled RGG condensates with 0 μM and 1.5 μM actin, displaying polymerized actin within the condensates and condensate deformation in the presence of actin. Scale bars, 5 μm.

